# Acetyl-Leucine slows disease progression in lysosomal storage disorders

**DOI:** 10.1101/2020.05.20.105973

**Authors:** Ecem Kaya, David A Smith, Claire Smith, Lauren Morris, Tatiana Bremova-Ertl, Mario Cortina-Borja, Paul Fineran, Karl J Morten, Joanna Poulton, Barry Boland, John Spencer, Michael Strupp, Frances M Platt

## Abstract

Acetyl-DL-leucine (ADLL) is a derivative of the branched chain amino acid leucine. In observational clinical studies ADLL improved symptoms of ataxia, in particular in patients with the lysosomal storage disorder (LSD), Niemann-Pick disease type C 1 (NPC1). Here, we investigated ADLL and its enantiomers acetyl-L-leucine (ALL) and acetyl-D-leucine (ADL) in symptomatic *Npc1*^*-/-*^ mice and observed an improvement in ataxia with both enantiomers and ADLL. When ADLL and ALL were administered pre-symptomatically to *Npc1*^*-/-*^ mice, both treatments delayed disease progression and extended life span, whereas ADL did not. These data are consistent with ALL being the neuroprotective enantiomer. Altered glucose and antioxidant metabolism were found to be implicated as one of the potential mechanisms of action of the L enantiomer in *Npc1*^*-/-*^ mice. When miglustat and ADLL were used in combination significant synergy resulted. In agreement with these pre-clinical data, when NPC1 patients were evaluated after 12 months of ADLL treatment, rates of disease progression were slowed, with stabilisation or improvement in multiple neurological domains. A beneficial effect of ADLL on gait was also observed in this study in a mouse model of GM2 gangliosidosis (Sandhoff disease) and in Tay-Sachs and Sandhoff disease patients in individual-cases of off-label-use. Taken together, we have identified an unanticipated neuroprotective effect of acetyl-L-leucine and underlying mechanisms of action in LSDs, supporting its further evaluation in clinical trials in lysosomal disorders.

## Introduction

Acetyl-DL-leucine (ADLL) is a racemic (1:1) mixture of enantiomers and a derivative of the branched chain amino acid leucine. It has been used as a medication (Tanganil™) for the treatment of acute vertigo in France and former French colonies since 1957; it is orally available and has a good safety profile (Neuzil *et al*., 2002). Although its mechanism of action is not fully known, studies in a guinea pig model of acute vertigo showed that ADLL can restore membrane potential and excitability of abnormally polarised neurons of the medial vestibular nucleus (Vibert and Vidal, 2001).

ADLL was shown to be beneficial in patients with cerebellar ataxias (Strupp *et al*., 2013; Schniepp *et al*., 2016). More recently, significant improvements in ataxia and fine motor function were documented in 12 patients with the rare lysosomal storage disorder (LSD), Niemann-Pick disease type C 1 (NPC1), who were treated with the oral formulation of ADLL (Bremova *et al*., 2015). Improvements in cognition and behaviour were also found, suggestive of potential functional benefits beyond the cerebellum (Bremova *et al*., 2015). Our findings on *in vitro* benefit of ADLL in Tangier disease fibroblasts (Colaco *et al*., 2019), and *in vivo* slowing of disease progression in Sandhoff disease mouse model (Kaya *et al*., 2020) suggesting ADLL may have the potential for therapeutic use in a broader range of diseases and neuroprotective effect.

In all of these studies, ADLL was administered to patients as a racemic mixture of acetyl-D-leucine (ADL) and acetyl-L-leucine (ALL) enantiomers. Pre-clinical studies implicated ALL as the active isomer responsible for postural compensation after unilateral vestibular damage in an animal model (Günther *et al*., 2015; Tighilet *et al*., 2015), and three clinical trials are ongoing with acetyl-L-leucine: NPC, GM2 gangliosidosis and ataxia telangiectasia (clinicaltrials.gov NCT03759639, NCT03759665, NCT03759678, and EudraCT (2018-004331-71; 018-004406-25; 2018-004407-39)). However, it remains to be determined if this proposed mechanism of action is relevant to the effects of ADLL and individual enantiomers in NPC.

In the current study, we have therefore investigated the efficacy of ADLL and its distinct enantiomer components in a mouse model of NPC1 (*Npc1*^*-/-*^). We found that ADLL and its distinct enantiomers significantly improved ataxia in *Npc1*^*-/-*^ mice, in agreement with clinical observations with ADLL (Bremova et al., 2015). It is important to note, when administered pre-symptomatically ADLL (50% ALL: 50% ADL) and ALL, but not ADL, slowed the rate of disease progression, were neuroprotective, and significantly extended lifespan. Therefore, pre-clinical studies implicate ALL as the active enantiomer of ADLL responsible for the neuroprotective effects of the treatment.

When miglustat, the currently approved therapy for NPC1 (Patterson *et al*., 2007), was combined with ADLL in *Npc1*^*-/-*^ mice, significant synergistic benefit resulted, including further extension of life span. In a cohort of NPC1 patients, the significant neuroprotective effects of ADLL identified in *Npc1*^*-/-*^ mice were also observed in a 12-18-month extension phase of an ongoing observational study (Cortina-Borja et al., 2018). When the mechanism of action of ADLL and its enantiomers were investigated in the cerebellum of treated *Npc1*^*-/-*^ mice, modulation of pathways involved in glucose metabolism were identified as potentially mediating the beneficial effects of the L-enantiomer. Finally, in individual-cases of off-label-use ADLL improved function in three GM2 gangliosidosis patients (Tay-Sachs and Sandhoff disease) and gait improvements were also demonstrated in a mouse model of Sandhoff disease. These findings demonstrate the distinct benefits of acetyl leucine treatments in LSDs, and show promise for clinical applications.

## Materials and Methods

**Animals and treatments**. BALBc/NPC1^nih^ (Pentchev *et al*., 1980) and Sandhoff (*Hexb*^-/-^) mice (Sango *et al*., 1995) were bred as heterozygotes to generate homozygous null (*Npc1*^*-/-*^) and mutant (*Hexb*^-/-^) mice, along with their respective wild-type controls (*Npc1*^*+/+*^, *Hexb*^*+/+*^). Mice were housed under non-sterile conditions, with food and water available *ad libitum*. All experiments were conducted on female mice to facilitate group housing from different litters using protocols approved by the UK Home Office Animal Scientific Procedures Act, 1986. All animal works was conducted under the UK Home Office licensing authority and the Project licence number is P8088558D. Mice were randomly assigned to treatment groups and the drugs coded and the staff blinded to treatment when performing behavioural analysis.

**Drug treatments**. Acetyl leucine analogues (ALs) and miglustat treatment protocols are described in the **supplemental experimental procedures**.

**Mouse behavioural analysis**. The weight and activity of each mouse was recorded weekly until reaching the humane endpoint (defined as a loss of 1g body weight within 24h). CatWalk (10.5 system (Noldus)), NG Rota Rod for mice (Ugo Basile) was performed as described in the **supplemental experimental procedures**.

### Biochemical and immunohistochemical analyses

**Sample preparation**. Mice were saline perfused under terminal anaesthesia. Tissues for biochemical analysis were snap-frozen on ice-cold isopentane. For immunofluorescent staining, tissues were perfused with 4% PFA followed by PBS and kept in 4% PFA for 24h then stored in PBS containing 20% w/v sucrose. Biochemical analyses were performed on water-homogenised tissues (50 mg/mL) and protein content determined (BCA protein assay, Thermo Fisher #23227) according to the manufacturer’s instructions.

**Western blot analyses**. Western blot analyses were performed on mouse cerebellum as described in the **supplemental experimental procedures**.

**ADP and ATP extraction**. ADP and ATP were extracted with Phenol-TE from tissues according to published methods (Chida *et. al*., 2012). For details, see **supplemental experimental procedures**.

**NAD and NADH extraction**. NAD and NADH extractions were performed on 20mg PBS washed fresh tissue on wet ice homogenised with a Dounce homogeniser with 400uL of NADH/NAD extraction buffer (Abcam NAD/NADH Assay kit-ab65348). For details, see **supplemental experimental procedures**.

### Sample analysis

**Sphingoid Base Measurements**. Sphingoid base extraction and detection with reverse phase HPLC was performed as described in the **supplemental experimental procedures**.

**Glycosphingolipid measurements**. Glycosphingolipids were extracted and measured by Normal Phase-HPLC according to published methods (Neville *et. al*., 2004). For details, see **supplemental experimental procedures**.

**Cholesterol measurements**. Cholesterol was measured from Folch-extracted tissues with the Amplex Red kit (Molecular Probes) according to the manufacturer’s instructions. For details, refer to **supplemental experimental procedures**.

**Flow cytometry experiments of CHO cells**. Relative acidic compartment volume staining of live cells was with LysoTracker™ Green DND-26 (Thermo Fisher-L7526), mitochondrial volume was determined with MitoTracker Green (Invitrogen #M7514) and mitochondrial reactive oxygen species with MitoSOX Red (Invitrogen #M36008). For details, refer to **supplemental experimental procedures**.

**Filipin Staining of CHO cells**. Free unesterified cholesterol was visualized in CHO cells grown on glass coverslips using filipin labelling (from Streptomyces filipinensis (Sigma)). For details, refer to **supplemental experimental procedures**.

**Immunohistochemistry**. Immunohistochemistry was performed as described in the **supplemental experimental procedures**. The antibodies used are summarised in **Supplementary Table 1**. Slides were air dried and mounted in ProLong Gold antifade (Invitrogen, #P36930).

**Table 1.** Number of patients (total cohort n=13) that improved, stabilised or deteriorated following ADLL treatment stratified by the neurological domains scored (NIH NCSS).

| **Neurological Domains** | **Improved** | **Stable** | **Deteriorated** |
| --- | --- | --- | --- |
| **Eye movement** | 8 | 4 | 1 |
| **Ambulation** | 8 | 4 | 1 |
| **Speech** | 3 | 7 | 3 |
| **Swallow** | 10 | 2 | 1 |
| **Fine motor skills** | 8 | 3 | 2 |
| **Cognition** | 6 | 6 | 1 |
| **Memory** | 10 | 2 | 1 |

**Western Blotting**. Western blotting was performed as described in **supplemental experimental procedures**. The antibodies used are summarised in **Supplementary Table 1**.

**Image Acquisition and Quantification**. Imaging of brain sections and CHO cells was performed with a Leica-SP8 confocal microscope. Western blot data acquisition was conducted with LiCOR Odyssey Infrared imaging system (Model No. 9120) and Universal hood II (Bio-Rad, California, USA). Mean fluorescence values, cerebellum diameters and areas, and cell quantifications were calculated with Fiji Version 1.51g software (Image J) (http://fiji.sc/Fiji) (W. Rasband, NIH, USA).

**ADP/ATP measurements**. Measurements of ADP/ATP were made using a kit from Sigma Aldrich (Cat#: MAK135) according to the manufacturer’s instructions. The ADP/ATP ratio was calculated with the equation: (RLU-C-RLUB)/RLU-A.

**NAD, NADH measurements**. NAD, NADH and total NAD (NADt) (NAD+NADH=NADt) were measured with the NAD/NADH assay kit (Abcam-ab65348). For details, refer to **supplemental experimental procedures**.

### Clinical Studies

***Demographics and statistical analysis of individual-cases of off-label-use of adult NPC1 patients treated with ADLL is*** described in the **supplemental experimental procedures**.

***Individual-cases of off-label-use in GM2 gangliosidosis patients*** are described in the

### supplemental experimental procedures

***Blinded Video-Rating*** is described in the **supplemental experimental procedures**.

All patients gave their informed consent according to the Declaration of Helsinki prior to the compassionate use study. The clinical examinations were approved by the ethical committee of Ludwig-Maximilians-University Munich.

### Statistical methods and Power calculations

To calculate the number of mice needed for the experimental groups in this study we used G*Power software (http://www.gpower.hhu.de). The power was set as 0.8 and the significance level, α, as 0.05. The mean lifespan of the NPC1 mouse model in our facility is 87 days with a standard deviation of 3 days. NPC1 is a disease with no curative treatment and many experimental disease-modifying drugs have reported a 10-30% increase in lifespan. Therefore, we based our power calculation on a 10-30% effect size and a sample size of min 5 and max 8 animals per group was determined. For statistical analysis see **supplemental experimental procedures**.

### Data availability

Data will be made available upon reasonable request.

## Results

### ADLL, ALL and ADL administered during the symptomatic phase of disease improves ataxia in Npc1^-/-^ mice

*Npc1*^*-/-*^ mice have a 10-12-week life span, with onset of symptoms (gait abnormalities, tremor and weight loss) beginning at 6-7 weeks of age (Williams *et al*., 2014). ADLL, ALL and ADL were administered to *Npc1*^*-/-*^ mice in their diet (0.1g/kg/day), with a dose identical to that used in observational clinical studies (Bremova *et al*., 2015). Untreated 9-week-old *Npc1*^*-/-*^ mice exhibit statistically significant ataxia that presents as a pronounced sigmoidal gait, relative to wild-type mice (*p*<0.0001) (**Suppl. Fig. 1**), whereas 9-week-old *Npc1*^*-/-*^ mice treated with ADLL, ALL or ADL from 8-weeks of age (1 week of treatment) all displayed significantly reduced ataxia as determined by measuring lateral displacement from a straight trajectory in an automated gait analysis system (**Suppl. Fig. 1**) **(Fig. 1a and b)** (*p*<0.0001, all treatments). These results indicate that the acute anti-ataxic effect of acetyl-leucine is stereoisomer-independent in *Npc1*^*-/-*^ mice.

**Figure 1.**
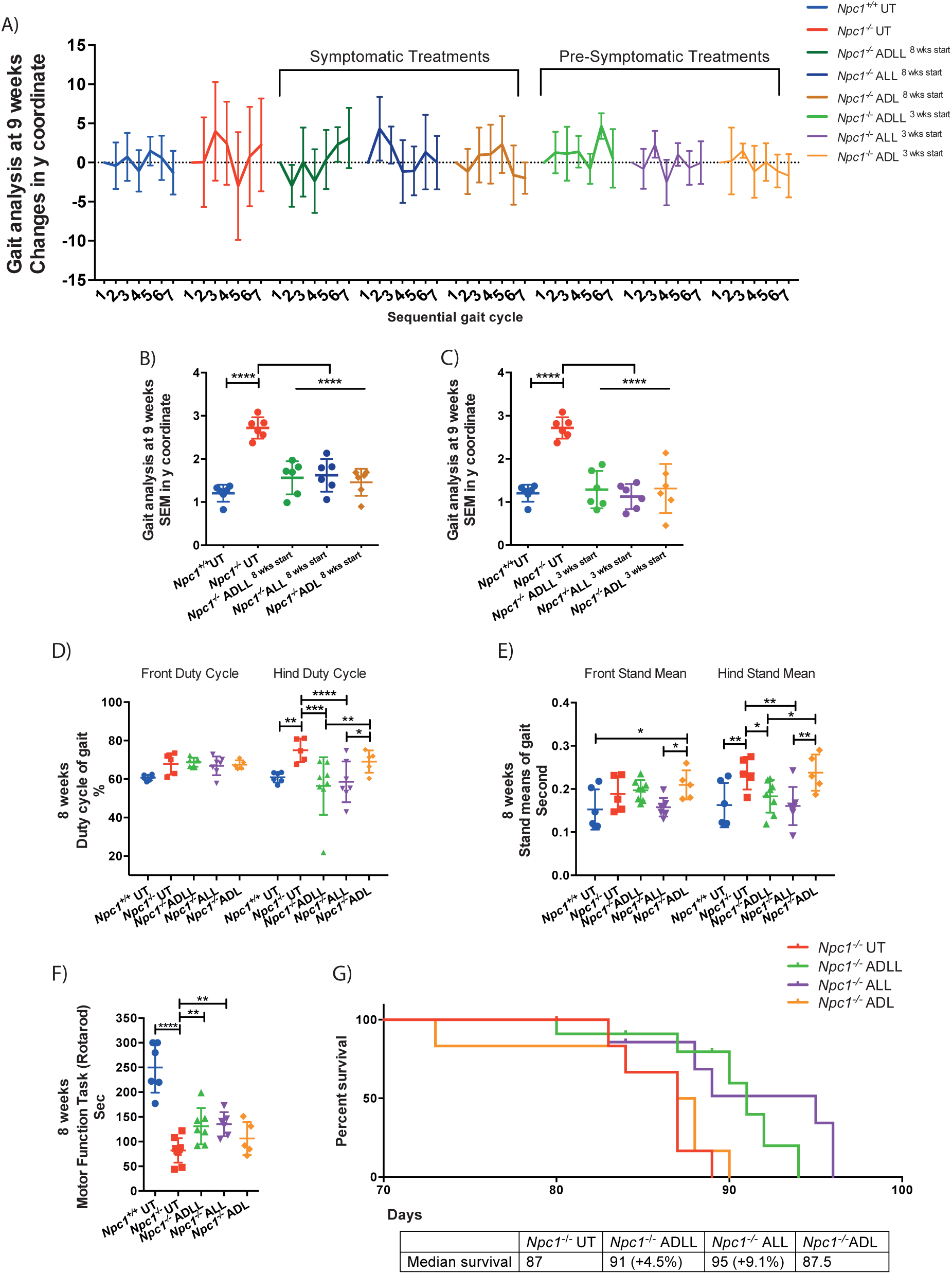
Acetyl leucine analogues impact on behavioural parameters in NPC1 mice. For wild type untreated (*Npc1*^*+/+*^ UT), NPC1 untreated (*Npc1*^*-/-*^ UT), ADLL (*Npc1*^*-/-*^ ADLL), ALL (*Npc1*^*-/-*^ ALL), ADL (*Npc1*^*-/-*^ ADL) treatments min 5 animals, max 7 for each group. **a)** Symptomatic and pre-symptomatic AL treatment Y coordinates displacement of each consecutive foot from a straight-line trajectory. (Mean ± SD, n=6) **b)** Late stage AL treatment SEMs of Y coordinate changes (Mean ± SD, n=6; *p<*0.0001, One-way ANOVA **c)** Early stage AL treatment SEMs of Y coordinate changes (Mean ± SD, n=6; *p<*0.0001, One-way ANOVA). **d)** Early stage AL treatments front and hind duty cycle measurements, mean ± SD, \*\**p*<0.032, \*\*\**p* <0.0005, \*\*\*\**p*<0.0001 (2-way ANOVA) **e)** Early stage AL treatments front and hind stand mean measurements, min 5 animals per group, Mean ± SD, * *p*<0.026, \*\**p*<0.003 (2-way ANOVA). **f)** Early stage AL treatment motor performance measurements, min 5 animals each group, Mean ± SD, \*\**p*<0.009, \*\*\*\**p*<0.0001 (One-way ANOVA). **g)** Life expectancy percentages, Mean ± SD, * *p*<0.046 (Gehan-Breslow-Wilcoxon test) n=6.

### Pre-symptomatic treatment with ADLL, ALL and ADL improves ataxia in Npc1^-/-^ mice

We investigated whether any additional benefit was conferred by AL analogues if treatment commenced before symptom onset. *Npc1*^*-/-*^ mice were therefore treated pre-symptomatically from weaning (3 weeks of age) and assessed at 9 weeks of age (6 weeks of treatment). Similar to the acute treatment response, all three AL analogues tested significantly reduced ataxia (**Fig. 1a and c**) (*p*<0.0001 for all treatments) with the magnitude comparable to the effects observed with one week of treatment (**Fig. 1a and Suppl. Fig. 1**).

### Pre-symptomatic treatment with ADLL and ALL, but not ADL, improves gait abnormalities, motor function and modestly extends survival in Npc1^-/-^mice

*Npc1*^*-/-*^ mice develop motor dysfunction (measured by Rotarod) and paw placement abnormalities when ambulatory (measured with CatWalk). We therefore treated *Npc1*^*-/-*^ mice pre-symptomatically from weaning (3 weeks of age) and assessed the mice at 8 weeks of age. Untreated *Npc1*^-/-^ mice had a significantly improved hind paw duty cycle (**Fig. 1d**, *p*=0.0031) and an improved hind paw stand mean (*p*=0.0026) relative to wild-type mice (Kirkegaard *et al*., 2016) **(Fig. 1e)**. Pre-symptomatic ADLL and ALL treatment of *Npc1*^*-/-*^ mice resulted in a functional improvement in duty cycle of hind paws (ADLL: *p*=0.0005, ALL: *p*<0.0001) **(Fig. 1d)** and hind paw stand mean (ADLL: *p*=0.0179, ALL: *p*=0.0014) relative to untreated *Npc1*^*-/-*^ mice **(Fig. 1e)**. Rotarod performance was significantly impaired in untreated *Npc1*^*-/-*^ mice relative to wild-type littermates (*p*<0.0001) (**Fig. 1f**). Treatment with ADLL (*p*=0.0088) and ALL (*p*=0.0070) resulted in a significant improvement in Rotarod performance by *Npc1*^*-/-*^ mice **(Fig. 1f)**. ADL treatment had no significant benefit on *Npc1*^*-/-*^ mouse motor function, as assessed using Catwalk (hind paw duty cycle *p*=0.2124, hind paw stand mean *p*=0.9547) or Rotarod (*p*=0.2218) **(Fig. 1d-f)**. Relative to untreated *Npc1*^*-/-*^ mice, the life span of animals treated from weaning was modestly but significantly increased by 8 days (9.1%) with ALL treatment (*p*=0.0334), 4 days with ADLL (*p*=0.0305) (4.5%) and was not changed with ADL (*p*=0.6908) **(Fig. 1g)**, therefore displaying L-enantiomer selectivity.

### ADLL, ALL and ADL improves neuropathology and lipid storage in Npc1^-/-^ mouse brain in a stereo-selective manner

Since NPC1 disease is characterized by the accumulation of sphingoid bases (sphingosine and sphinganine), cholesterol, sphingomyelin, free fatty acids and glycosphingolipids (GSLs) (Lloyd-Evans and Platt, 2010) we measured the impact of ALs administered to *Npc1*^*-/-*^ mice from 3 weeks of age on lipid storage in the brain. At 59 days of age *Npc1*^*-/-*^ mice exhibited increased lipid levels relative to wild-type (Williams *et al*., 2014) (**Fig. 2a-f**). In order to evaluate the differential impact of the AL analogues in the CNS we analyzed the cerebellum and the forebrain (referred to as brain in the **Fig. 2**) separately. Total GSLs in the forebrain were not significantly altered by any of the AL treatments **(Suppl. Fig. 2a)**, whilst ALL selectively reduced GM1a (20.1%; *p=*0.0018) and GM2 (19.6%; *p=*0.0222) **(Fig. 2a)**. Interestingly, although sphingosine levels were not significantly affected, ADLL and ADL reduced sphinganine levels by 13.5% (*p=*0.0456) and 18.2% (*p=*0.0111) respectively **(Fig. 2b and c)**. Analysis of the cerebellum confirmed no significant difference in total GSLs relative to untreated *Npc1*^*-/-*^ mice with any of the AL analogues tested **(Suppl. Fig. 2b)**. However, ADLL and ALL treatment significantly lowered levels of specific GSLs including GA2 (ADLL: 21.3%, *p=*0.0102; ALL: 23.8%, *p*=0.0042), whereas a significant reduction in GA1 was only observed with ALL treatment (23.5%, *p=*0.0254) **(Fig. 2d)**. Comparable to the results for the forebrain, sphingosine was unchanged in the cerebellum, but sphinganine was significantly reduced following ADL treatment (42.3%, *p=*0.0074) **(Fig. 2e and f)**. These results indicate differential biological targets of the two enantiomers.

**Figure 2.**
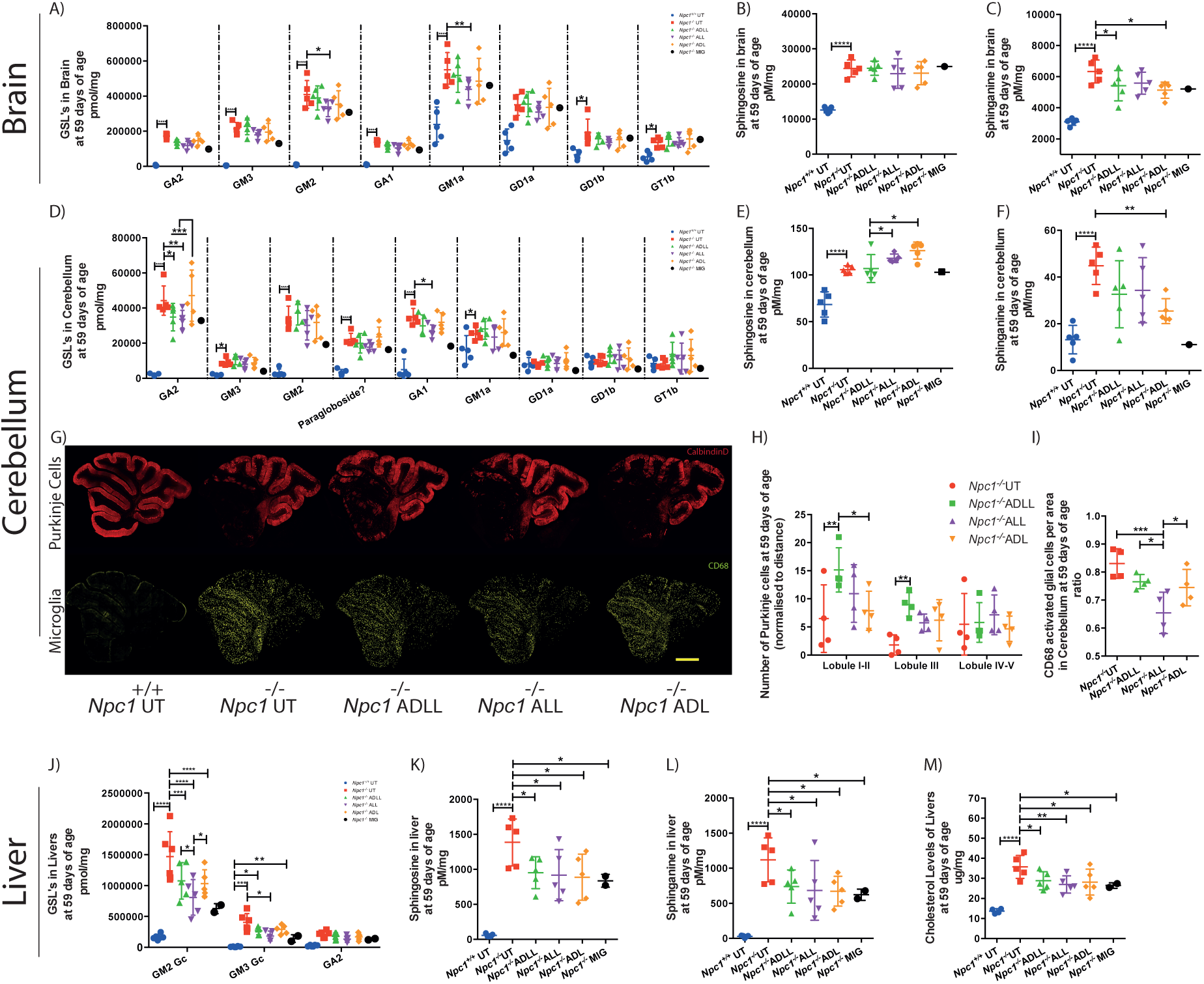
Efficacy of Acetyl leucine analogues on biochemical and histopathology in *Npc1*^*-/-*^ mice at 59 days of age. For all AL treatments n=5 animals per group, for miglustat (positive control) (*Npc1*^*-/-*^ MIG) n=2 or n=1. **a)** Glycosphingolipid (GSL) profiles in brain, mean ± SD, \**p*<0.024, \*\**p*<0.0019, \*\*\**p*<0.0005, \*\*\*\**p*<0.0001(2-way ANOVA). **b)** Sphingosine levels in brain, mean ± SD, \*\*\*\**p*<0.0001(One-way ANOVA). **c)** Sphinganine levels in brain, mean ± SD, \**p*<0.046, \*\*\*\**p*<0.0001(One-way ANOVA). **d)** GSLs in cerebellum, mean ± SD, \**p*<0.026, \*\**p*<0.005, \*\*\**p*<0.0009, \*\*\*\**p*<0.0001 (2-way ANOVA). **e)** Sphingosine in cerebellum, mean ± SD, * *p*<0.046\*\*\*\**p*<0.0001 (One-way ANOVA). **f)** Sphinganine in cerebellum, mean ± SD, ** *p*<0.075, \*\*\*\**p*<0.0001(One-way ANOVA). **g)** Cerebellum stained with calbindin-b (Purkinje cells, red) or CD68 (activated microglia, yellow). Scale bar 800 um. **h)** Purkinje cell density at 59 days of age relative to NPC1 untreated, mean ± SD, * *p*<0.011, ** *p*<0.028 (One-way ANOVA) **i)** CD68 cell density at 59 days of age, mean ± SD, * *p*<0.046, *** *p*<0.0009 (One-way ANOVA). **j)** GSLs in liver, mean ± SD \**p*<0.038, *** *p* <0.0002, \*\*\*\**p*<0.0001 (2-way ANOVA). **k)** Sphingosine in liver, mean ± SD. \**p*<0.0393, \*\*\*\**p*<0.0001(One-way ANOVA) **l)** Sphinganine in liver, mean ± SD, \**p*<0.04, \*\*\*\**p*<0.0001(One-way ANOVA). **m)** Liver cholesterol, mean ± SD, \**p*<0.034, \*\**p*=0.0091, \*\*\*\**p*<0.0001(One-way ANOVA).

### ADLL slows Purkinje cell loss and ALL reduces neuroinflammation in the cerebellum, whereas ADL has no neuroprotective effect

*Npc1*^*-/-*^ mice exhibit progressive neurodegeneration, with cerebellar Purkinje neuron loss progressing from anterior to posterior lobes, accompanied by microglial activation (Williams *et al*., 2014) (**Fig. 2g**). Only ADLL treatment significantly increased Purkinje cell survival at 59 days of age relative to untreated *Npc1*^*-/-*^ littermates: 133% more Purkinje cells were present in lobules I and II (*p=*0.0027), and 402% more Purkinje cells in lobule III (*p=*0.0108) (**Fig. 2g and h)**. Other treatments did not significantly improve Purkinje cell survival in any cerebellar lobules (*Npc1*^*-/-*^ALL lobule I-II *p=*0.107, lobule III *p=*0.157, lobule IV-V *p=*0.533; *Npc1*^*-/-*^ADL lobule I-II *p=*0.60, lobule III *p=*0.11, lobule IV-V *p=*0.766, compared to *Npc1*^*-/-*^ UT) (**Fig. 2g and h)**. In addition, only ALL treatment significantly reduced (by 20%, *p=*0.0177) the frequency of CD68-positive activated microglia **(Fig. 2g and i)** while other treatments did not have a significant impact (*p=0*.*1353 for Npc1*^*-/-*^ADLL, *p=0*.*0553 for Npc1*^*-/-*^ADL compared to *Npc1*^*-/-*^ UT). ADL did not mediate any long term, neuroprotective effects assessed with Purkinje cell count and CD68 staining.

### All AL treatments alleviate lipid storage in non-neuronal tissues and cells

Liver GSL levels in the *Npc1*^*-/-*^ mice were significantly reduced in *Npc1*^*-/-*^ mice treated with ADLL (26.9%, *p=*0.03), ADL (26.9%, *p=*0.0253) and 45.5% for ALL (*p=*0.0003) (**Suppl. Fig. 2c**). Quantification of the major GSL species confirmed that levels of GM2Gc (the most abundant GSL in the mouse liver) was decreased significantly with all AL analogues tested (26.7% ADLL: *p=*0.0002, 45% ALL: *p<*0.0001, 29% ADL: *p<*0.0001), while GM3Gc levels were only reduced significantly by ALL (54.8%: *p=*0.0314) relative to untreated *Npc1*^*-/-*^ mice **(Fig. 2j)**. All analogues tested caused significant and comparable levels of reduction of sphingosine (the catabolic break down product of ceramide) (ADLL 29.5%, *p=*0.0233; ALL 33.2%, *p*=0.0148; ADL 33.6% *p*=0.0103) and its *de novo* precursor sphinganine (ADLL 32.5% *p=*0.0365; ALL 37.4% *p*=0.0189; ADL 38.5% *p*=0.0163) **(Fig. 2k and l)**. Total free cholesterol in *Npc1*^*-/-*^ liver was also significantly decreased after treatment with ADLL, ALL or ADL (ADLL 18.2% *p=*0.0346; ALL 23.4% *p*=0.0091; ADL 16.2 % *p*=0.0210) **(Fig. 2m)**.

Reduction of lipid species in the liver with all three AL treatments led us to investigate whether other non-neuronal cell types were corrected by ALs. We conducted a series of experiments on Npc1 deficient Chinese Hamster Ovary (CHO) cells by utilizing lysosomal and mitochondrial probes as well as measurements of lipid storage. NPC1 protein deletion/mutation results in increased lysosomal volume (Te Vruchte *et al*., 2014), as well as an increase in mitochondria (measured by MitoTracker), mitochondrial superoxide increase (measured by MitoSOX) (Paulina Ordonez *et al*., 2012). CHO cells treated with 1mM ALs for 24 hours resulted in normalisation of MitoSOX staining with all three drugs (*p<*0.0001, compared to NPC1 UT) **(Fig. 3a)**, and a significant decrease in MitoTracker Green (ADLL 16.8% *p*=0.0150; ALL 17.1% *p*=0.0133; ADL 15.8% *p*=0.0215) **(Fig. 3a)**. We found that only 1mM ADLL and ALL treatment reduced relative lysosomal volume after 24h (ADLL 16.9%, *p=*0.0137; ALL 16%, *p*=0.0177) **(Fig. 3a)**; however, this beneficial effect was seen with all three acetyl leucine treatments after 72h of treatment (ADLL 24% *p=*0.0026; ALL 22.8% *p*=0.0036; ADL 22.3 % *p*=0.0042) **(Fig. 3a)**. After 72 hr AL treatment, sphingosine levels were significantly reduced by 22% with ADLL (*p*=0.0009), 29% with ALL (*p*<0.0001) and 14% with ADL (*p*=0.0149) **(Fig. 3b)**. Total GSL levels were significantly reduced with all treatments (26% with ADLL *p*=0.0007; 25% with ALL *p*=0.0010; and 22% with ADL *p*=0.0024) **(Fig. 3b)**. Likewise, cholesterol content was significantly reduced (18% with ADLL *p*= 0.0043; 13% with ALL *p*=0.0263; and 14% with ADL *p*=0.0212) **(Fig. 3b, c)**.

**Figure 3.**
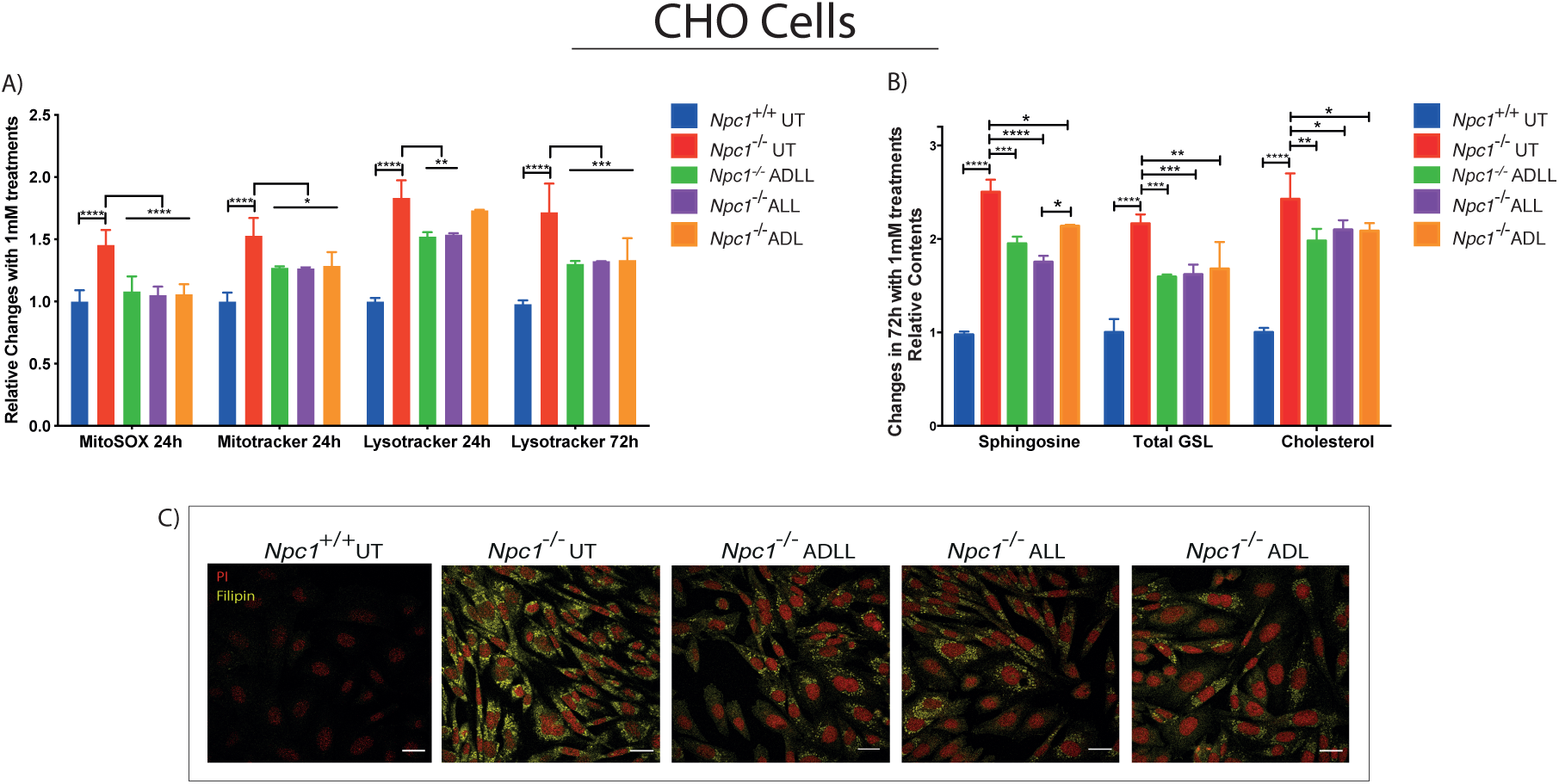
Efficacy of Acetyl leucine analogues on NPC1-deficient CHO cells. **a)** MitoSOX(red) and Mitotracker(Green) and Lysotracker(Green) changes relative to *Npc1*^+/+^ with 24 and 72h 1mM ALs Mean ± SD, \**p*<0.022, \*\**p*<0.0063, \*\*\**p*<0.0007, **** *p*<0.0001 (2-way ANOVA). **b)** Sphingosine, total GSL and free cholesterol levels (72h) relative to *Npc1*^+/+^, Mean ± SD, \**p*<0.027, \*\**p*<0.0044, \*\*\**p*<0.001 \*\*\*\**p*<0.0001 (2-way ANOVA). **c)** NPC1 CHO cells stained with filipin (yellow, cholesterol) and propidium iodide (red, nucleus). Scale bar 20 micron.

### ADLL synergises with miglustat in Npc1^-/-^ mice

We treated *Npc1*^*-/-*^ mice with ADLL in combination with the standard of care drug miglustat (600mg/kg/day), resulting in a statistically significant longer life span than animals on monotherapy (*Npc1*^*-/-*^ miglustat vs *Npc1*^*-/-*^ miglustat & ADLL *p=*0.0016; *Npc1*^*-/-*^ ADLL vs *Npc1*^*-/-*^ miglustat & ADLL *p=*0.0063; median survival; *Npc1*^*-/-*^ UT: 87 days, *Npc1*^*-/-*^ ADLL: 91 days; *Npc1*^*-/-*^ miglustat: 117 days, *Npc1*^*-/-*^ miglustat & ADLL: 138 days) **(Fig. 4a)**. Miglustat is known to cause weight loss through appetite suppression (Priestman *et al*., 2008). When ADLL was used in combination with miglustat it prevented the weight loss associated with miglustat treatment **(Fig. 4b)** but not the weight loss resulting from progression of the disease. Combination therapy significantly increased Rotarod performance (8 weeks: *p=*0.0007; 10,12 weeks *p<0*.*0001* versus untreated *Npc1*^*-/-*^), with combination therapy associated with increased benefit relative to mice receiving either miglustat (10 weeks: *p=*0.0299; 12 weeks: *p*=0.0232) or ADLL monotherapy (10,12 weeks: *p<0*.*0001*) **(Fig. 4c)**. Gait analysis at 10 weeks of age demonstrated that hind Duty Cycle percentages were significantly improved with combination therapy (*p=*0.0028) whereas miglustat monotherapy showed no benefit **(Fig. 4d and e)**. Stand mean significantly improved in mice treated either with combination therapy (*p=*0.0250) or miglustat monotherapy (*p=*0.0196) **(Fig. 4d, f)**. Due to limited availability of miglustat, ALL and ADL combination therapies were not evaluated.

**Figure 4.**
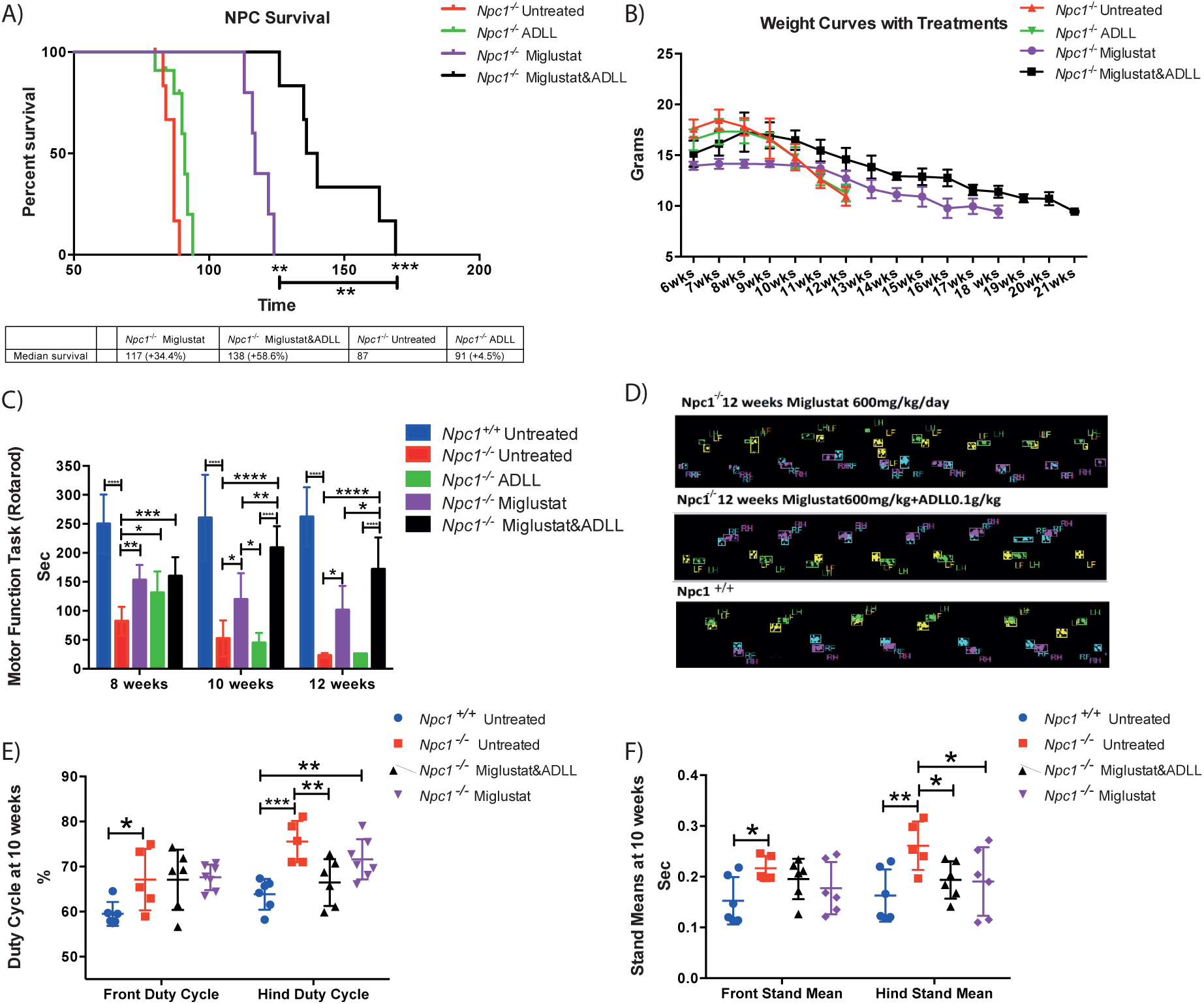
Effect of ADLL in combination with miglustat. For wild type untreated (*Npc1*^*+/+*^ UT), NPC1 untreated (*Npc1*^*-/-*^ UT), ADLL (*Npc1*^*-/-*^ ADLL), miglustat (*Npc1*^*-/-*^ MIG) and combination (*Npc1*^*-/-*^ miglustat & ADLL) therapies min 5 animals for each group. Duty Cycle and Stand mean analyses were measured by 3 recorded runs per animal and quantitative results were obtained with Noldus CatWalk 10.5 software system. Motor function performance was measured with accelerating Rotarod (1rpm per minute up to 10 rpm). **a)** Survival curves, Mean ± SD, \**p*=0.0270, \*\**p*<0.0035, \*\*\**p*=0.0007 (Gehan-Breslow-Wilcoxon test). **b)** Body weight curves over time, Mean ± SD. **c)** Motor performance measurements at 8, 10 and 12 weeks of age, Mean ± SD, \**p*<0.043, \*\**p*<0.0036, \*\*\*\**p*< 0.0001 (2-way ANOVA) **d)** Representative image of footprints of miglustat and miglustat/ADLL combination therapy relative to wild type **e)** Duty cycle measurements of front and hind paws at 10 weeks of age, Mean ± SD, \**p*=0.011, \*\**p*<0.0053, \*\*\**p* 0.0002 (2-way ANOVA) **f)** Stand mean measurements at of front and hind paws 10 weeks of age, Mean ± SD, \**p*<0.033, \*\**p*<0.0016 (2-way ANOVA).

### Potential targets of Leucine analogues in the cerebellum

We investigated the potential targets that ALs could affect in the cerebellum since this brain region is particularly important in NPC1-related pathology. As L-Leucine is a potent activator of the mammalian target of rapamycin (mTOR) (Kimball *et al*., 1999), which negatively regulates autophagy (Klionsky and Emr, 2000), we investigated mTOR and its phosphorylation status in the cerebellum of AL treated *Npc1*^*-/-*^ *mice*. Western blotting indicated that levels of mTOR and its phosphorylation on serine 2448 (Yanagisawa *et al*., 2017) (phosphorylated/total protein expression ratios (p/t)) were not changed by ADLL, ALL or ADL **(Suppl. Fig. 3a and b)**. Autophagic vacuoles accumulate in NPC1 disease, consistent with impaired lysosomal flux (Boland *et al*., 2010). None of the compounds significantly changed the ratio of LC3-I and LC3-II **(Suppl. Fig. 3b and c)**. Furthermore, none of the ALs changed autophagic function or flux **(Suppl. Fig. 3b and c)**. In addition, we were unable to detect a significant change in levels of its substrate p62/SQSTM1(Pankiv *et al*., 2007) **(Suppl. Fig. 3b and d)**.

#### Effects on energy metabolism and anti-oxidant pathways

In order to investigate potential disruption of branched chain amino acid metabolism we examined mitochondrial branched chain keto acid dehydrogenase enzyme A subunit (BCKADHA), which functions in the final step of the pathway that yields acetyl CoA (Harris *et al*., 2005). Phosphorylation of BCKADHA (that inactivates the enzyme) was significantly reduced in untreated *Npc1*^*-/-*^ cerebellum relative to wild type (37.6%, *p=*0.0254) and therefore enzyme activity (predicted by the ratio of phosphorylated enzyme (p) and total enzyme (t)-p/t) was significantly increased (36.6%, *p=*0.0269) **(Suppl. Fig. 3e and f)** in *Npc1*^*-/-*^ cerebellum. None of the ALs tested had a significant impact on either total BCKADHA levels or the extent of its phosphorylation **(Suppl. Fig. 3e and f)**. Branched chain amino acids, especially leucine, are important for glutamine/glutamate balance and for energy metabolism when glucose metabolism is deficient or impaired (Yudkoff, 1997) (Kennedy *et al*., 2016). However, there was no difference in mitochondrial enzyme glutamate dehydrogenase (GDH) protein levels between untreated *Npc1*^*-/-*^ and *Npc1*^*+/+*^ mice or following AL treatments **(Suppl. Fig. 3f and g)**.

The status of glucose metabolism can be assessed by determining adenosine diphosphate and adenosine triphosphate ratios (ADP/ATP), and the ratio of nicotinamide adenine dinucleotide and its reduced form (NAD/NADH). We therefore measured the ADP/ATP and NAD/NADH ratios. NAD and NADH were decreased in *Npc1*^*-/-*^ relative to *Npc1*^*+/+*^ mice achieving statistical significance for NADH (55.1%, *p=*0.0314) and sum of NAD and NADH (49.2%, *p=*0.0001) **(Fig. 5a)**. Although there was no change in the levels of the coenzymes following AL treatments, in comparison with untreated NPC1 **(Fig. 5a)**, the NAD/NADH ratio was significantly decreased with ADLL treatment (30.7%, *p=*0.0240) **(Fig. 5b)**, which would be indicative of increased glycolysis. In accordance with these data, the ADP/ATP ratio was also increased with ADLL **(Fig. 5c)** (*p=*0.0101).

**Figure 5.**
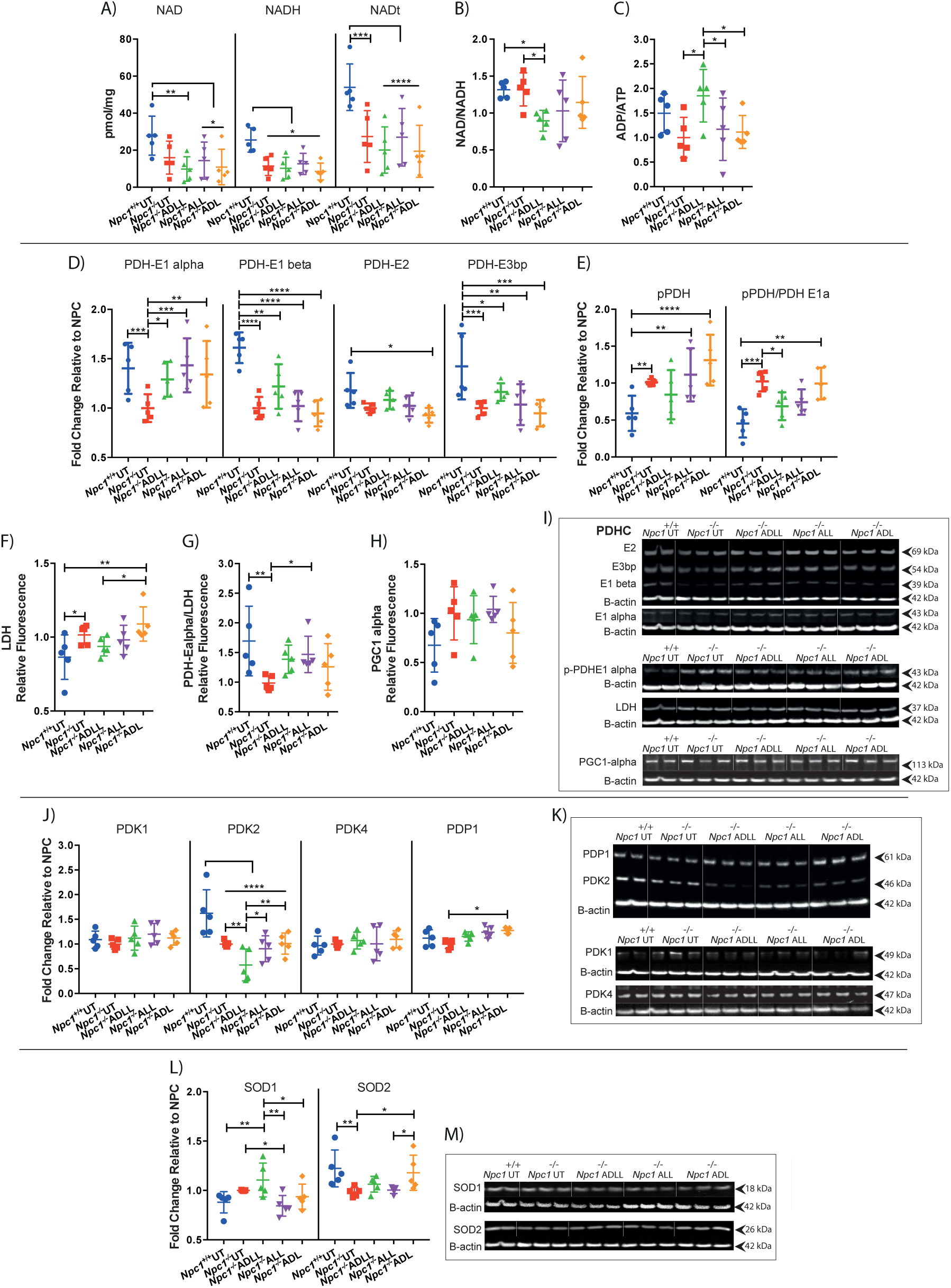
Effects of acetyl leucine analogues on expression of proteins involved in energy metabolism and the antioxidant system in cerebellum at 59 days of age. Wild type untreated (*Npc1*^*+/+*^ UT), NPC1 untreated (*Npc1*^*-/-*^ UT), ADLL (*Npc1*^*-/-*^ ADLL), ALL (*Npc1*^*-/-*^ ALL), ADL (*Npc1*^*-/-*^ ADL) treatments, n=5 for each group. **a)** NAD, NADH, total NAD (NADt) measurements, mean ± SD, \**p*<0.05, \*\**p*=0.0065, \*\*\**p*=0.0001, \*\*\*\**p*<0.0001 (2-way ANOVA). **b)** NADH/NADH ratios, mean ± SD, \**p*<0.025 (One-way ANOVA). **c)** ADP/ATP ratios, Mean ± SD (One-way ANOVA). **d)** PDHC protein expressions, Mean ± SD, \**p*<0.034, \*\**p*<0.0045, \*\*\**p*<0.001, \*\*\*\**p*<0.0001 (2-way ANOVA). **e)** pPDH expressions and pPDH/PDHE1 alpha ratios, Mean ± SD, \**p*=0.0333, \*\**p*<0.0095, \*\*\**p*=0.0006, \*\*\*\**p*<0.0001 (2-way ANOVA). **f)** LDH expression, Mean ± SD, \**p*<0.034, \*\**p*<0.0028 025 (One-way ANOVA). **g)** PDHE1 alpha/LDH ratios, Mean ± SD, \**p*=0.0494, \*\**p*=0.0060 (One-way ANOVA). **h)** PGC 1 alpha expression, mean ± SD (One-way ANOVA). **i)** Western blot images of PDHC, pPDH, LDH, PGC-1alpha and n-actin loading controls. **j)** PDK 1-2-4 and PDP1 expression, Mean ± SD, \**p*<0.045, \*\**p*<0.0023, \*\*\*\**p*<0.0001 (2-way ANOVA). **k)** Western blot images of PDKs and PDP1 along with their n-actin loading controls. **l)** SOD1 and SOD2 expression, Mean ± SD, \**p*<0.0472 \*\**p*<0.0064 (2-way ANOVA). **m)** Western blot images of SOD1 and SOD2 along with beta-actin loading controls.

#### ADLL increases active pyruvate dehydrogenase (PDH) levels while ALL shifts glucose metabolism towards PDH from lactate dehydrogenase (LDH)

We investigated the effect of AL analogues on glucose utilization, by measuring levels of pyruvate dehydrogenase (PDH) and lactate dehydrogenase (LDH) proteins by western blotting **(Fig. 5d-i)**. PDH is composed of 3 subunits: E1-3, with the active site located on the alpha subunit of E1. Enzyme activity is reduced by phosphorylation(Kato *et al*., 2008). Untreated *Npc1*^-/-^ mice at 59 days of age showed significantly decreased levels of PDH-E1 alpha (40.2%, *p=*0.0009), PDH-E1 beta (61.2%, *p<*0.0001), PDH-E3 binding protein (PDH-E3bp) (42.2%, *p=*0.0005) **(Fig. 5d and i)**, while phosphorylation of PDH (41.4%, *p=*0.0092), PDH enzyme activity (pPDH/PDH E1alpha) (55.5%, *p=*0.0006) and LDH increased in the disease group (14.2%, *p=*0.0335) **(Fig. 5e, f and i)**, in comparison to WT. Accordingly, the glucose metabolism preference (PDHE1 alpha/LDH) favored LDH dependency in untreated *Npc1*^-/-^ mice (74.2%, *p=*0.0060) with no detectable difference in proliferator-activated receptor-gamma coactivator 1-alpha levels **(Fig. 5 g-i)**. E1 alpha subunit protein levels in PDHC were significantly increased by ADLL (29.1%, *p*=0.0146), ALL (43.3%, *p*=0.0004) and ADL (34.17%, *p*=0.0044) in the cerebellum from treated *Npc1*^*-/-*^ mice **(Fig. 5d and i)**, with no changes in El beta, E2 or E3bp. The protein levels of phosphorylated E1 alpha subunit were not significantly different following treatment compared to untreated *Npc1*^*-/-*^ mice. However, the ratio of phosphorylated E1 alpha to total E1 alpha subunit was reduced with ADLL (33%, *p*=0.0333) **(Fig. 5e and i)**. LDH levels were the same in untreated and treated *Npc1*^*-/-*^ mice **(Fig. 5f and i)**. Nevertheless, the ratio of E1 alpha subunit to LDH was significantly shifted in favor of PDH by ALL (49.1%, *p*=0.0494) **(Fig. 5g)**. Finally, we measured the protein levels of PGC-1 alpha (peroxisome proliferator-activated receptor-gamma coactivator 1-alpha), a key regulator of oxidative phosphorylation (Huiyun and Walter, 2006), to determine whether this metabolic activation and shift towards PDH dependency had an impact on oxidative phosphorylation. There was no difference in protein levels of PGC-1 alpha between WT and NPC1 cerebellum, or associated with AL treatments **(Fig. 5h and i)**.

#### ADLL decreases pyruvate dehydrogenase kinase 2 (PDK2) protein expression

Pyruvate dehydrogenase is phosphorylated and inactivated by pyruvate dehydrogenase kinase (PDK), reducing flux through the Krebs cycle. In contrast, pyruvate dehydrogenase phosphatase (PDP) dephosphorylates and activates PDH, increasing flux through the Krebs cycle (Jha *et al*., 2012). We therefore analyzed the expression of three PDK isoforms (PDK1, PDK2 and PDK4) and a single isoform of PDP (PDP1) in the cerebellum (Jha *et al*., 2012). Whilst levels of PDK1 and PDK4 in the cerebellum were not significantly different between 59-day old *Npc1*^*-/-*^ and *Npc1*^*+/+*^mice, PDK2 was significantly lower (64.4% reduction compared to wild type, *p<*0.0001) **(Fig. 5j and k)**. Furthermore, ADLL (which decreased the phosphorylation of PDH to the greatest extent) decreased PDK2 to a significantly greater extent relative to *Npc1*^*-/-*^ UT group (42.3%, *p=*0.0022) **(Fig. 5j and k)**. PDP1 was unaltered in untreated *Npc1*^*-/-*^ cerebellum and was only significantly increased by ADL (27.4%, *p=*0.0433) **(Fig. 5j and k)**.

#### Differential effects of individual enantiomers on antioxidant systems: ALL decreases superoxide dismutase (SOD) 1 while ADL increases SOD2

We assessed the impact of ALs on the anti-oxidative system by measuring the cytoplasmic superoxide dismutase 1 (SOD1) and the mitochondrial superoxide detoxifier superoxide dismutase 2 (SOD2). We were unable to detect any significant difference in SOD1 protein levels between *Npc1*^*-/-*^ and *Npc1*^*+/+*^ cerebella but observed a significant reduction in SOD2 (25% *p*=0.0061) **(Fig. 5l and m)**. We found that only ALL significantly decreased SOD1 levels in cerebellum (15.6%, *p=*0.0472) and only ADL significantly elevated SOD2 levels when compared to untreated *Npc1*^*-/-*^ mice (19.8%, *p*=0.0214) **(Fig. 5l and m)**.

### Neuroprotective effects of ADLL in individual-cases of off-label-use in NPC1 patients

Having observed unanticipated beneficial effects of ALs in *Npc1*^*-/-*^ mice we investigated whether neuroprotective effects also occurred in NPC1 patients treated with ADLL enrolled in an observational clinical study (Cortina-Borja *et al*., 2018). Total clinical severity scores (with higher values equating to increasing levels of disability (Yanjanin *et al*., 2010)) were plotted prior to initiation of treatment with Tanganil™ (ADLL), incorporating available retrospective data (**Fig. 6a**). All 13 patients had positive slopes of disease progression before ADLL treatment (1.8 severity units/year was the average slope). Following initiation of ADLL treatment the average slope was -1.78 (3 patients had no slope, 10 had negative slopes) equating to a significant reduction in disease progression and improvement in the majority of patients (One-sided sign test *p*= 0.0002) (**Fig. 6a**). The data were also computed as Annual Severity Increment Scores (Cortina-Borja *et al*., 2018) that measures the rate of disease progression. A mean of -9.1% / year (*p*<0.001) in ASIS scores was observed **(Fig. 6b)**. When the slope each patient’s pre-treatment total severity score per year was plotted, all patients had positive slopes, whereas post treatment slopes were either zero or negative, consistent with stabilisation or improvement in disease progression **(Fig. 6c)**. When the treated patient data were analysed by neurological subdomains (**Table 1**) the majority of patients improved or stabilised on treatment in functional and cognitive subdomains **(Table 1)**.

**Figure 6.**
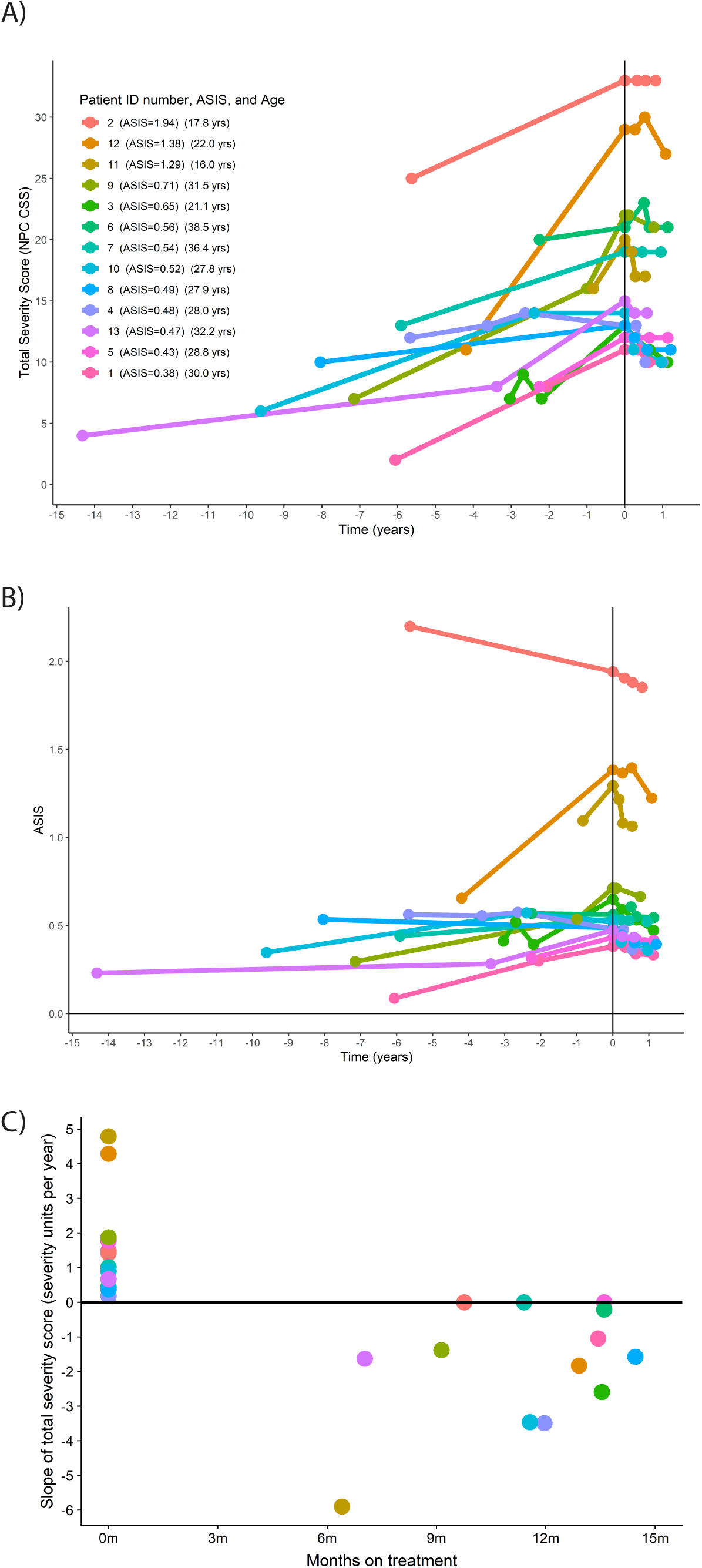
Clinical data from 13 adult NPC1 patients. For demographics see materials and methods **a)** Total clinical severity scores (maximum score 50, with higher values equating to increasing levels of disability(Yanjanin *et al*., 2010)). Data were plotted prior to initiation of treatment with Tanganil™ (ADLL), incorporating available retrospective data **b)** Clinical data computed as Annual Severity Increment Scores (Cortina-Borja *et al*., 2018), (*p*<0.001 in ASIS score) **c)** Slope of total severity score (severity units per year) before and after ADLL treatment. Each individual participant in this observational study is colour coded.

### ADLL shows benefit in other LSDs: Effects in Sandhoff mice and GM2 gangliosidosis patients

The Sandhoff (*Hexb*^*-/-*^) mouse model is pre-symptomatic up to 6-8 weeks of age. Subsequently, they develop tremor and their motor function begins to decline (Jeyakumar *et al*., 1999). By the later stages of the disease (12-15 weeks) they are inactive and are unable to complete motor function tests such as bar crossing (Jeyakumar *et al*., 1999). *Hexb*^*-/-*^ animals were treated from weaning with ADLL (0.1mg/kg/day, the same dose used for treatment of *Npc1*^*-/-*^ mice). ADLL-treated mice had improved gait parameters; including hind stand mean (*p=*0.0323) **(Fig. 7a)**, front (*p=*0.0039) and hind step cycle (*p=*0.0062) **(Fig. 7b)**. To date, ALL and ADL have not been evaluated in the Sandhoff mouse model.

**Figure 7.**
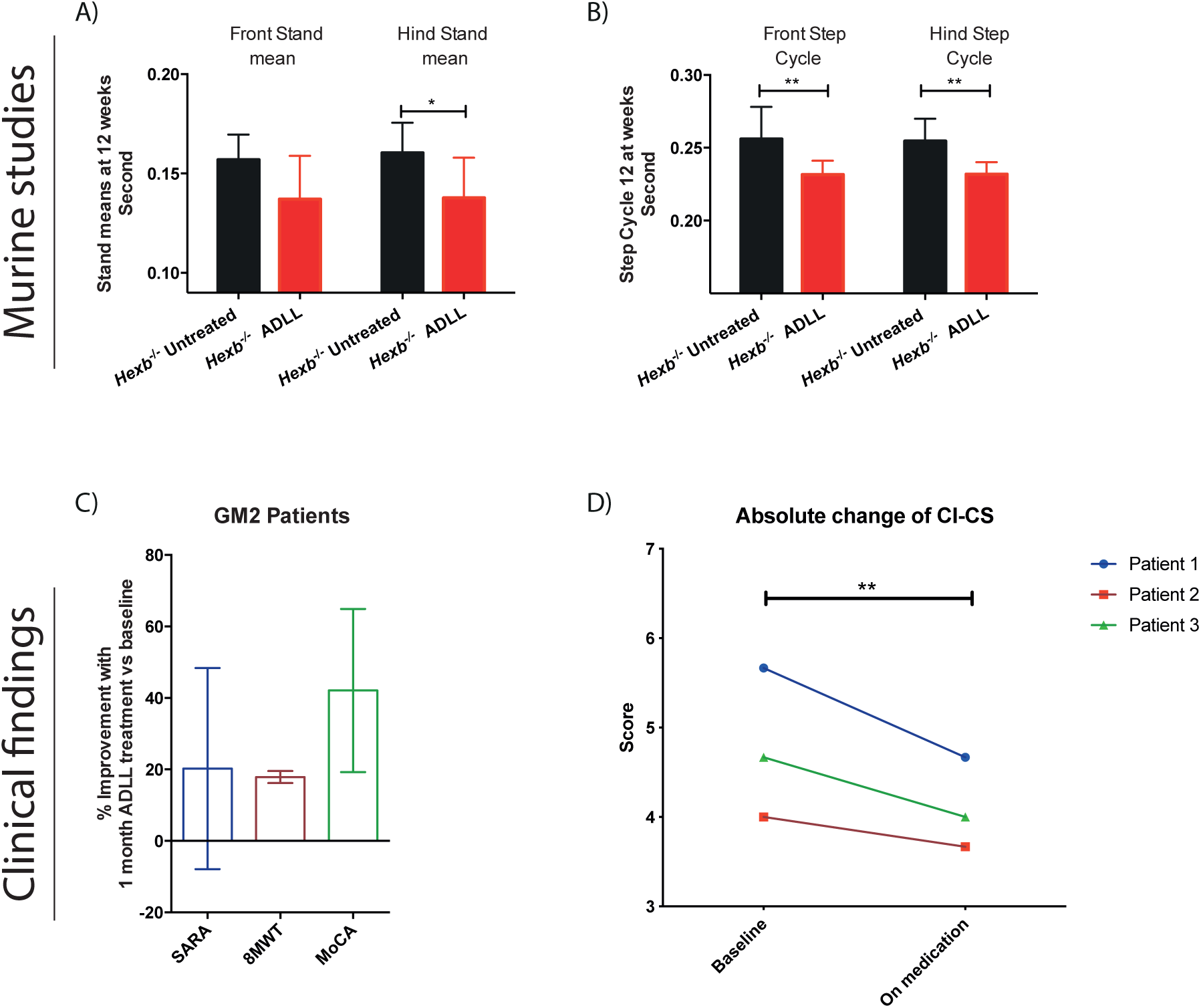
Effects of Acetyl-DL-Leucine on behavioural and biochemical parameters in 12-week-old Sandhoff disease mice. Sandhoff untreated (*Hexb*^*-/-*^ UT), ADLL (*Hexb*^*-/-*^ ADLL). 5 animals per group. **a)** Front-hind stand mean measurements of *Hexb*^*-/-*^ UT and *Hexb*^*-/-*^ADLL, Mean ± SD, \**p*=0.0323, 2-way ANOVAs **b)** Front-hind step cycle measurements of *Hexb*^*-/-*^ UT and *Hexb*^*-/-*^ADLL, Mean ± SD, \*\**p<*0.0063 2-way ANOVAs **c)** Percent improvement in clinical scores in three patients with GM2 gangliosidosis on baseline and after one month on medication with N-Acetyl-DL-Leucine (ADLL). Regarding the motor scores, Scale for Assessment and Rating of Ataxia (SARA) changed by 20.3% and the 8-M-Walking-Test (8MWT) yielded a change by 17.8%. Cognitive function, as assessed by Montreal Cognitive Assessment (MoCA) changed by 42.0%. See also **Supplementary videos 1-2. d)** Unbiased Clinical Impression of Change in Severity (CI-CS) scores of GM2 patient videos. CI-CS Skala Clinical Impression of Change in Severity; 1=normal, not at all ill; 2=borderline ill; 3=mildly ill; 4=moderately ill; 5=markedly ill; 6=severely ill; 7=among the most extremely ill patients. Paired t test, p=0.0039. For individual scoring data see **Table 2**.

Our findings in Sandhoff mice were extended to individual-cases of off-label-use in three patients with a confirmed GM2 gangliosidosis diagnosis (2 Tay-Sachs and 1 Sandhoff disease (the latter case in press (Bremova-Ertl *et al*., 2020)) treated with ADLL. We found a mean improvement of the Scale for Assessment and Rating of Ataxia (SARA) by 20.3%, the 8-M-Walking-Test (8MWT) by 17.8% and the Montreal Cognitive Assessment (MoCA) by 42%, as shown in **Fig, 7c**. All patients and caregivers also reported a subjective improvement and have continued treatment at the same dosage. Videos of the effect of treatment on gait and postural instability are shown in **Supplemental Videos 1-2**. In addition, three experienced movement disorder experts performed blinded analysis of the videos, and rated the videos based on the Clinical Impression of Severity (CI-S) (1=normal, not at all ill; 2=borderline ill; 3=mildly ill; 4=moderately ill; 5=markedly ill; 6=severely ill; 7=among the most extremely ill). The unbiased observation before and after ADLL treatment showed statistically significant improvements on overall scoring (t-test *p*=0.0039) **(Fig. 7d and Table 2)**. ALL and ADL have not been evaluated in clinical settings, but trials with ALL are being conducted (NPC (NCT03759639) and GM2 gangliosidosis (NCT03759665) and Ataxia-Telangiectasia (NCT03759678)).

**Table 2.** **CI-CS scoring of 3 GM2 patients by 3 blinded and unbiased experts**. Blinded scoring of GM2 gangliosidosis patient videos pre and post ADLL treatment. CI-CS Skala Clinical Impression of Change in Severity 1=normal, not at all ill; 2=borderline ill; 3=mildly ill; 4=moderately ill; 5=markedly ill; 6=severely ill; 7=among the most extremely ill patients

| **Examiner Number** | **Patient 1 Baseline** | **Patient 1 Treated** | **Patient 2 Baseline** | **Patient 2 Treated** | **Patient 3 Baseline** | **Patient 3 Treated** | **Paired t-test** |
| --- | --- | --- | --- | --- | --- | --- | --- |
| **1** | 5 | 4 | 3 | 3 | 3 | 3 |  |
| **2** | 6 | 5 | 4 | 3 | 5 | 4 |  |
| **3** | 6 | 5 | 5 | 5 | 6 | 5 |  |
| **Mean** | 5.67 | 4.67 | 4 | 3.67 | 4.67 | 4 | *p*=0.0039 |

## Discussion

In this study we have investigated the effects of ADLL, ALL and ADL in mouse models of NPC1, Sandhoff disease and in patients with NPC1 patients and GM2 gangliosidosis to better understand the therapeutic potential of these drugs and to gain insights into their mechanisms of action.

We found that ADLL, ALL and ADL significantly improved ataxia when symptomatic *Npc1*^*-/-*^ mice were treated acutely for seven days; this is in agreement with observational studies in NPC1 patients (Bremova *et al*., 2015) treated with ADLL, using the same dosage per kg and day. The individual enantiomers (ALL and ADL) provided similar benefit to the racemic mixture (ADLL) for the symptomatic treatment of ataxia. The mechanism(s) explaining how ADLL and both the D and L enantiomers rapidly improve symptoms of ataxia remains unknown. All AL analogues tested reduced lipid storage in neuronal and non-neuronal tissues in *Npc1*^*-/-*^ mice and in CHO cells null for *Npc1* suggesting a mechanism of action that is not confined to neuronal cells as previously suggested (Vibert and Vidal, 2001). The AL analogues were also observed to differentially reduce the levels of stored lipids in the liver and, to a lesser extent, in the brain of treated *Npc1*^*-/-*^ mice. The mechanism that underpins this “substrate reduction” action of ALs currently remains unclear, but in view of the high degree of synergy when combined with the *bona fide* substrate reduction therapy drug miglustat, it may not be a major contributor to AL’s therapeutic effect in NPC1 disease.

The major finding of the current study was that ADLL and ALL (but not ADL) slowed disease progression when treatment was initiated before symptom onset, consistent with a neuroprotective mechanism. Since the neuroprotective effects were only observed with ADLL and ALL, this implicated ALL as the active enantiomer and demonstrates that the symptomatic improvement in ataxia and neuroprotection are achieved through different mechanisms of action. ALL significantly reduced neuroinflammation, which is important as CNS inflammation actively contributes to disease progression and reducing inflammation using nonsteroidal anti-inflammatory drugs has previously been shown to be beneficial in *Npc1*^*-/-*^ mice (Williams *et al*., 2014).

The neuroprotective effect of ADLL and ALL prompted us to determine whether similar effects occur in NPC1 patients treated with ADLL. We took advantage of an ongoing observational study in which thirteen NPC1 patients had been treated with Tanganil™ (ADLL) continuously for approximately 1 year and found that all patients showed stabilisation or improvement in clinical scores following treatment. Significantly, these improvements were across all neurological domains, not just those relating to ataxia, supporting a more global neuroprotective effect in patients, analogous to those observed in the *Npc1*^*-/-*^ mouse.

One central question arising from these studies is the nature of the underlying mechanism(s) through which ADLL and its separate enantiomers provide symptomatic, and in the case of ADLL and ALL, neuroprotective benefit in NPC1. Therefore, we investigated aspects of cell biology and metabolism known to be sensitive to leucine. Leucine has been shown to activate mTOR and reduce LC3-II and p62 in NPC1 cells (Yanagisawa *et al*., 2017). However, in this study, ADLL, ALL and ADL did not significantly affect either mTOR or autophagy in *Npc1*^*-/-*^ mouse cerebellum. This might be due to the presence of the acetamide group, as opposed to a primary amine in leucine, blocking the interaction with mTOR, recently shown in Hela cells (Nagamori *et al*., 2016) or that acetyl leucine analogues distribute at the cellular level in a manner that prevents their interaction with mTOR. Catabolism of leucine serves as a source of Acetyl-CoA (Harris *et al*., 1985) that has been shown to activate mTOR and nutrient sensing pathways (Son *et al*., 2018). However, the NPC1 cerebellum has significantly decreased levels of acetyl-CoA (Kennedy *et al*., 2016). This low acetyl-CoA content in NPC1 might therefore prevent nutrient sensing pathway activation. The complexity of mTOR pathways and altered metabolism more generally in NPC1 makes it likely that the effects of AL will likely be context specific.

Altered glucose/energy metabolism has previously been documented in the pre-symptomatic *Npc1*^*-/-*^ mouse cerebellum (Kennedy *et al*., 2013). PDHE1 alpha was found to be inactivated and there was a shift from pyruvate/PDH dependency (upon which neurons rely for aerobic respiration) towards lactate/lactate dehydrogenase (i.e. anaerobic respiration). The consequence of this would be progressive impairment of energy generation (Kennedy *et al*., 2013). In this same study, alterations in cerebellar amino acids (including leucine), and low acetyl CoA were also reported (Kennedy *et al*., 2013). Independent metabolomics studies on *Npc1*^*-/-*^ mouse liver reported imbalances in amino acid levels (Ruiz-Rodado *et al*., 2016). Interestingly, leucine was elevated in *Npc1*^-/-^ mice at both 3 and 5 weeks of age (along with branched chain amino-acid transferase 1) in forebrain and cerebellum, which could be a compensatory response for defective pyruvate oxidation and subsequent deficient energy production in models of NPC1 disease (Kennedy *et al*., 2016).

In view of data implicating altered energy metabolism in *Npc1*^*-/-*^ mice, we investigated the effects of ALs on glucose metabolism in the cerebellum of treated mice. We measured ADP/ATP and NAD/NADH ratios, as a sensitive measure of energy status (Liana Roberts Stein and Imai, 2012; Murphy and Hartley, 2018) and found that ADLL treatment shifted the system towards a more glycolytic state, in the absence of significant changes to NAD-NADH coenzyme levels, (which are lower in *Npc1*^*-/-*^ cerebellum), in agreement with previous findings in *Npc1*^*-/-*^ liver (Ruiz-Rodado *et al*., 2016). We then studied whether the shift in glycolysis is towards pyruvate utilization to produce lactate (anaerobic) or to produce acetyl CoA (aerobic). We found that ADLL treatment significantly increasing PDHE1alpha levels, and decreased PDH enzyme inactivation (phosphorylation) in the cerebellum. Interestingly, PDH-deficient mice also display Purkinje neuron degeneration and relapsing ataxia (Pliss *et al*., 2013), suggesting the ability of ADLL to protect Purkinje cells against degeneration may be via activation of PDH i.e. boosting aerobic (**Fig. 5**). This effect on PDH did not reflect a change in PGC1-alpha protein levels (indicator of oxidative phosphorylation, OXPHOS), which might be due to the fact that up-regulating OXPHOS requires coenzyme NADH and ADLL treatment does not increase NADH levels.

Individual enantiomers had distinct effects in NPC1 cerebellum; while ALL normalizes altered levels of PDH and LDH and mildly reduces SOD1 levels, ADL enhanced levels of the mitochondrial ROS scavenger SOD2 (manganese-dependent SOD2). *In vivo*, ADLL and ALL treatments were associated with a significantly greater benefit in motor function and lifespan relative to ADL **(Fig. 1)**, consistent with a possible connection between energy metabolism and motor function in NPC1. This study highlights that targeting altered cellular metabolism in neurodegenerative diseases may achieve greater efficacy than using anti-oxidant approaches. The metabolic changes in *Npc1*^*-/-*^ mice in response to AL treatments are summarized in **Fig. 8**.

**Figure 8.**
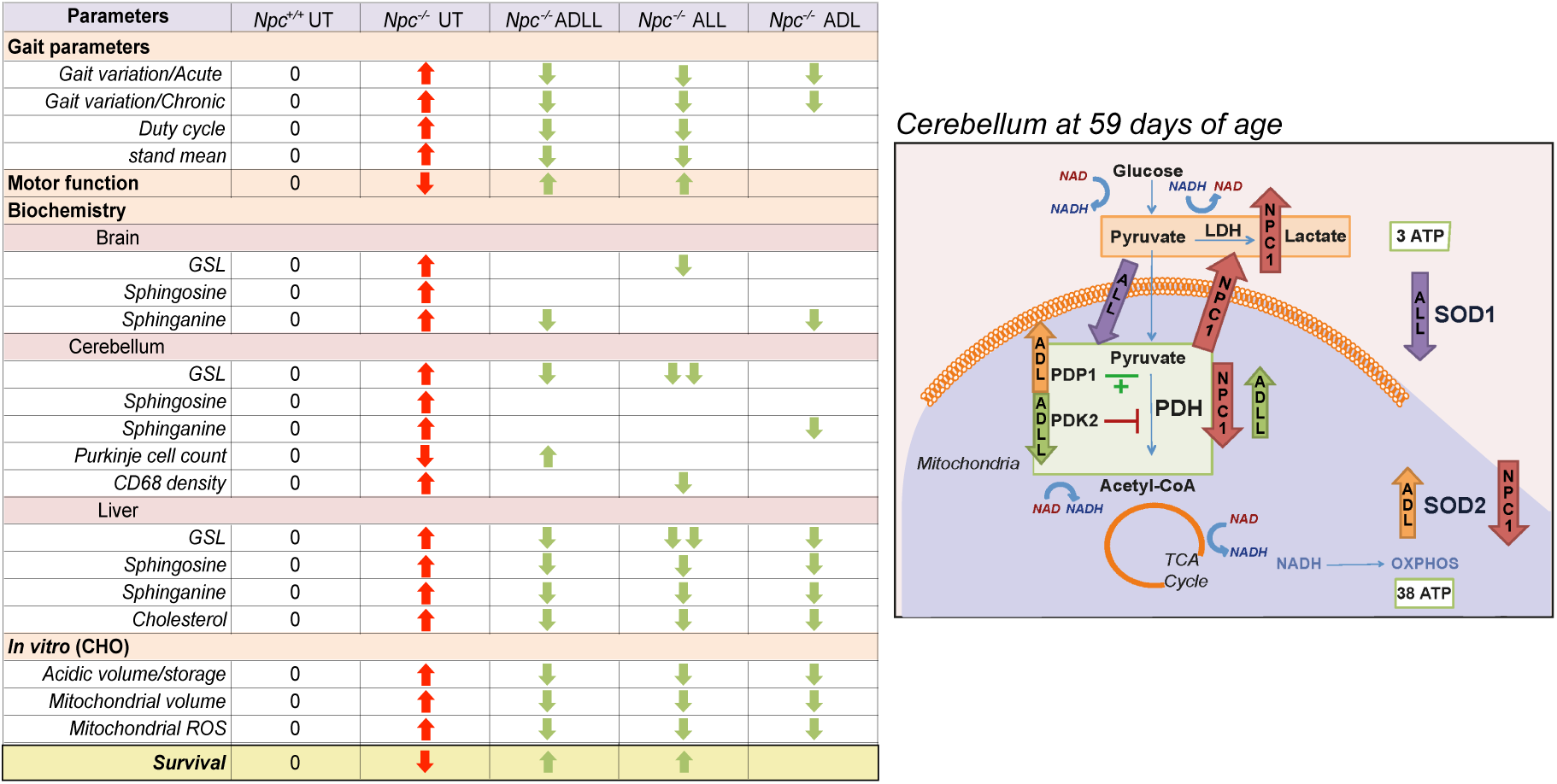
Summary table. Summary table of the effect of AL treatments on investigated pathologies *in vivo, ex vivo* and *in vitro*, along with the proposed mechanism of AL treatments in the cerebellum. Red arrows indicate the pathologies in NPC1 relative to its wild-type counterparts. Green arrows indicate the significant changes in response to acetyl-leucine treatment.

Finally, we explored whether the benefits of AL are confined to NPC1 or extend to other LSDs. We observed efficacy of ADLL in a mouse model of Sandhoff disease (the individual isomers ALL and ADL have not been assessed in the Sandhoff disease mouse model). Sandhoff disease is one of the GM2 gangliosidoses (Platt *et al*., 2012), a subgroup of GSL lysosomal storage diseases that includes Tay-Sachs disease, Sandhoff disease and GM2 activator deficiency. Sandhoff disease is caused by mutations in *HEXB*, a gene that encodes the beta subunit of β-hexosaminidase, leading to storage of GM2 ganglioside and other substrates in the CNS, leading to progressive neurodegeneration and premature death (Wiederschain, 2002). Sandhoff disease is associated with several phenotypes (developmental regression, profound intellectual disability, seizures and GSL accumulation) associated with deficits in energy production (Sandhoff and Harzer, 2013). We recently demonstrated that ADLL treated Sandhoff mice had a significant increase in lifespan and improvements in motor function (Kaya *et al*., 2020). In this study, we show improvement in gait parameters in *Hexb*^*-/-*^ mice upon ADLL therapy consistent with the observational clinical studies we conducted. Three patients with GM2 gangliosidosis with ataxia that were treated with ADLL in blinded individual-cases of off-label-use showed significantly improved gait, supporting the concept that ADLL and ALL may also be efficacious in multiple LSDs and other neurodegenerative diseases.

In conclusion, we have found that a well-tolerated drug, ADLL, and its enantiomer ALL slow disease progression and improve motor function and lipid accumulation in murine and cell models of NPC1. Furthermore, in observational studies in NPC1 patients, treatment with ADLL was associated with improvement in multiple neurological domains and a significant reduction in the rate of disease progression, potentially via restoration of aerobic metabolism based on the mechanistic studies conducted in the *Npc1*^*-/-*^ mouse model. The striking synergy of ADLL with miglustat in *Npc1*^*-/-*^ mice suggests that combining these two drugs will lead to improved short- and long-term clinical outcomes and merits clinical trials. It is interesting to note that 12 of the 13 NPC1 patients in the observational clinical study were on the standard of care miglustat, so part of their clinical improvement may be the result of the synergy between the two disease modifying drugs. The individual enantiomers ALL and ADL have not yet been tested in combination with miglustat. The finding that both a mouse model of Sandhoff disease and GM2 gangliosides patients, regardless of age of onset, also showed clinical improvement when treated with ADLL suggest that this, and related analogues, may have broader utility in LSDs and more common neurodegenerative diseases. Based on ALL being the neuroprotective enantiomer, clinical trials with ALL are currently being conducted (NPC (NCT03759639), GM2 gangliosidosis (NCT03759665) and Ataxia-Telangiectasia (NCT03759678)).

## Acknowledgements

We thank Ines Debove, MD, Joan Michelis, MD and Susanne Schneider, MD, PhD for blinded scoring of the GM2 gangliosidosis patient videos.

## Abbreviations

ADLL: Acetyl-DL-Leucine
NPC1: Niemann-Pick Disease Type C 1
LSD: Lysosomal Storage Diseases
ALL: Acetyl-L-Leucine
ADL: Acetyl-D-Leucine
GSL: Glycosphingolipid
CHO: Chinese Hamster Ovary
mTOR: Mammalian Target of Rapamycin
BCKADHA: Branched Chain Keto Acid Dehydrogenase Subunit A
GDH: Glutamate Dehydrogenase
ADP: Adenosine Diphosphate
ATP: Adenosine Triphosphate
NAD: Nicotinamide adenine dinucleotide
NADH: Nicotinamide adenine dinucleotide (reduced form)
PDH: Pyruvate Dehydrogenase
LDH: Lactate Dehydrogenase
PGC-1alpha: Peroxisome Proliferator-activated Receptor-gamma Coactivator 1-alpha
PDP: Pyruvate Dehydrogenase Phosphatase
PDK: Pyruvate Dehydrogenase Kinase
SOD: Superoxide Dismutase
ASIS: Annual Severity Increment Scores
SARA: Scale for Assessment and Rating of Ataxia
8MWT: 8-Minute-Walking-Test
MoCA: Montreal Cognitive Assessment
CI-CS: Clinical Impression of Change in Severity

## Figure Legends

**Supplementary Figure 1. Gait of mice with AL treatments initiated in the symptomatic or pre-symptomatic period**. Heat maps of mice which display the body route and footprints was obtained by Noldus Catwalk software. Red indicates the heat of the mouse body.

**Supplementary Figure 2. Total Glycosphingolipids in brain, cerebellum and liver at 59 days of age**. For wild type untreated (*Npc1*^*+/+*^ UT), NPC1 untreated (*Npc1*^*-/-*^ UT), ADLL (*Npc1*^*-/-*^ ADLL), ALL (*Npc1*^*-/-*^ ALL), ADL (*Npc1*^*-/-*^ ADL) treatments n=5 animals per group, miglustat treatment (*Npc1*^*-/-*^ MIG) n=2 or n=1. **a)** Total GSL measurements in forebrain (brain), Mean ± SD, \*\*\*\**p*<0.0001 (2-way ANOVA). **b)** Total GSL measurements in cerebellum, Mean ± SD, \*\*\*\**p*<0.0001 (2-way ANOVA). **c)** Total GSL measurements in liver, Mean ± SD, \**p*<0.034, \*\*\**p*=0.0004, \*\*\*\**p*<0.0001 (2-way ANOVA).

**Supplementary Figure 3. The effects of Acetyl leucine analogues on autophagy, BCKADH and glutamate dehydrogenase in NPC1 cerebellum at 59 days of age. a)** mTOR, Phospho-mTOR expressions and ratios between phospho-mTOR and mTOR, Mean ± SD (One-way ANOVA). **b)** Western blots images of phospho-mTOR, mTOR, LC3 and p62 and their beta actin loading controls. **c)** LC3-I, LC3-II expressions, and ratio of LC3-II versus LC3-I. Mean ± SD \*\*\*\**p*<0.0001 (2-way ANOVA). **d)** p62 levels, Mean ± SD, \*\**p*<0.006 (One-way ANOVA). In each experimental group n=5 mice **e)** BCKADH, pPBCKADH, p/t BCKADH expression, mean ± SD, \**p*<0.027, \*\**p*<0.005 (2-way ANOVA). **f)** BCKADH, pPCKADH, and GDH western blot images with beta-actin loading control. **g)** GDH expression, mean ± SD (One-way ANOVA).

**Supplementary Table 1**. List of antibodies /reagents used for immunohistochemistry and western blotting.

**Supplementary Video 1**

A 28-year old male patient with juvenile-onset Tay-Sachs disease, Eight-Meter Walk Test is shown. Note the gait instability with increased fall risk at baseline. The patient is not able to walk independently, constant support of either assistant, or the wall is required. After one month on medication with N-Acetyl-DL-Leucine (ADLL), the patient has more postural stability, the gait pattern changes with increased pace rate and reduced variability and postural sway.

**Supplementary Video 2**

Eight-Meter-Walk-Test of a 32-year old female patient with adult-onset Tay-Sachs disease is shown prior to and after 1 month on medication with N-Acetyl-DL-Leucine (ADLL). Note the more stable nature of the gait pattern with decreased fall risk on medication.

## References

Boland B, Smith DA, Mooney D, Jung SS, Walsh DM, Platt FM. Macroautophagy is not directly involved in the metabolism of amyloid precursor protein. J Biol Chem 2010; 285: 37415–37426.

Bremova T, Malinova V, Amraoui Y, Mengel E, Reinke J, Kolnikova M, et al. Acetyl-DL-leucine in Niemann-Pick type C: A case series. Neurology 2015; 85: 1368–1375.

Bremova-Ertl T, Platt FM, Strupp M. Sandhoff-disease: improvement of gait by acetyl-DL-leucine– a case report. Neuropediatrics (in press)

Colaco A, Kaya E, Adriaenssens E, Davis LC, Zampieri S, Fernández-Suárez ME, et al. Mechanistic convergence and shared therapeutic targets in Niemann-Pick disease type C and Tangier disease. J Inherit Metab Dis 2019: 1–12.

Cortina-Borja M, Te Vruchte D, Mengel E, Amraoui Y, Imrie J, Jones SA, et al. Annual severity increment score as a tool for stratifying patients with Niemann-Pick disease type C and for recruitment to clinical trials. Orphanet J Rare Dis 2018; 13: 1–16.

Günther L, Beck R, Xiong G, Potschka H, Jahn K, Bartenstein P, et al. N-Acetyl-L-Leucine accelerates vestibular compensation after unilateral labyrinthectomy by action in the cerebellum and thalamus. PLoS One 2015; 10: 1–18.

Harris A, Paxton R, Powell SM, Gillim SE. Physiological Covalent Regulation of Rat Liver Branched-Chain a-Ketoacid Dehydrogenase. Arch Biochem Biophys 1985; 243: 542–555.

Harris R a, Joshi M, Jeoung NH, Obayashi M. Overview of the molecular and biochemical basis of branched-chain amino acid catabolism. J Nutr 2005; 135: 1527S–30S.

Huiyun L, Walter WF. PGC-1alpha: a key regulator of energy metabolism. Adv Physiol Educ 2006; 30: 145–151.

Jeyakumar M, Butters TD, Cortina-Borja M, Hunnam V, Proia RL, Perry VH, et al. Delayed symptom onset and increased life expectancy in Sandhoff disease mice treated with N-butyldeoxynojirimycin. Proc Natl Acad Sci U S A 1999; 96: 6388–93.

Jha MK, Jeon S, Suk K. Pyruvate Dehydrogenase Kinases in the Nervous System: Their Principal Functions in Neuronal-glial Metabolic Interaction and Neuro-metabolic Disorders. Curr Neuropharmacol 2012; 10: 393–403.

Kato M, Wynn RM, Chuang JL, Tso SC, Machius M, Li J, et al. Structural Basis for Inactivation of the Human Pyruvate Dehydrogenase Complex by Phosphorylation: Role of Disordered Phosphorylation Loops. Structure 2008; 16: 1849–1859.

Kaya E, Smith DA, Smith C, Boland B, Strupp M, Platt FM. Beneficial Effects of Acetyl-DL-Leucine (ADLL) in a Mouse Model of Sandhoff Disease. [Internet]. J Clin Med 2020; 9 Available from: http://www.ncbi.nlm.nih.gov/pubmed/32276303

Kennedy BE, Hundert AS, Goguen D, Weaver ICG, Karten B. Presymptomatic Alterations in Amino Acid Metabolism and DNA Methylation in the Cerebellum of a Murine Model of Niemann-Pick Type C Disease. Am J Pathol 2016; 186: 1–18.

Kennedy BE, LeBlanc VG, Mailman TM, Fice D, Burton I, Karakach TK, et al. Pre-symptomatic activation of antioxidant responses and alterations in glucose and pyruvate metabolism in niemann-pick type C1-deficient murine brain. PLoS One 2013; 8: e82685.

Kimball SR, Shantz LM, Horetsky RL, Jefferson LS. Leucine regulates translation of specific mRNAs in L6 myoblasts through mTOR-mediated changes in availability of eIF4E and phosphorylation of ribosomal protein S6. J Biol Chem 1999; 17: 11647–52.

Kirkegaard T, Gray J, Priestman DA, Wallom KL, Atkins J, Olsen OD, et al. Heat shock protein-based therapy as a potential candidate for treating the sphingolipidoses. Sci Transl Med 2016; 8

Klionsky DJ, Emr SD. Autophagy as a regulated pathway of cellular degradation. Science 2000; 290: 1717–1721.

Liana Roberts Stein, Imai S. The dynamic regulation of NAD metabolism in mitochondria. Trends Endocrinol Metab 2012; 23: 420–428.

Lloyd-Evans E, Platt FM. Lipids on trial: The search for the offending metabolite in Niemann-Pick type C disease. Traffic 2010; 11: 419–428.

Murphy MP, Hartley RC. Mitochondria as a therapeutic target for common pathologies. Nat Rev Drug Discov 2018; 1: 865–886.

Nagamori S, Wiriyasermkul P, Okuda S, Kojima N, Hari Y, Kiyonaka S, et al. Structure-activity relations of leucine derivatives reveal critical moieties for cellular uptake and activation of mTORC1-mediated signaling. Amino Acids 2016; 48: 1045–1058.

Neuzil E, Ravaine S, Cousse H. La N-acetyl-DL-leucine, medicament symptomatique des etats vertigineux. Bull Soc Pharm Bordeaux 2002; 141: 15–38.

Pankiv S, Clausen TH, Lamark T, Brech A, Bruun JA, Outzen H, et al. p62/SQSTM1 binds directly to Atg8/LC3 to facilitate degradation of ubiquitinated protein aggregates by autophagy. J Biol Chem 2007; 282: 24131–45.

Patterson MC, Vecchio D, Prady H, Abel L, Wraith JE. Miglustat for treatment of Niemann-Pick C disease: a randomised controlled study. Lancet Neurol 2007; 6: 765–772.

Paulina Ordonez M, Roberts EA, Kidwell CU, Yuan SH, Plaisted WC, Goldstein LSB, et al. Disruption and therapeutic rescue of autophagy in a human neuronal model of Niemann pick type C1. Hum Mol Genet 2012; 21: 2651–2662.

Pentchev PG, Gal AE, Booth AD, Omodeo-Sale F, Fours J, Neumeyer BA, et al. A lysosomal storage disorder in mice characterized by a dual deficiency of sphingomyelinase and glucocerebrosidase. Biochim Biophys Acta - Lipids Lipid Metab 1980; 3: 669–79.

Platt FM, Boland B, van der Spoel AC. The cell biology of disease: lysosomal storage disorders: the cellular impact of lysosomal dysfunction. J Cell Biol 2012; 199: 723–34.

Pliss L, Hausknecht KA, Stachowiak MK, Dlugos CA, Richards JB, Patel MS. Cerebral Developmental Abnormalities in a Mouse with Systemic Pyruvate Dehydrogenase Deficiency. PLoS One 2013; 8: 1–14.

Priestman DA, Van Der Spoel AC, Butters TD, Dwek RA, Platt FM. N-butyldeoxynojirimycin causes weight loss as a result of appetite suppression in lean and obese mice. Diabetes, Obes Metab 2008; 2: 159–66.

Ruiz-Rodado V, Nicoli ER, Probert F, Smith DA, Morris L, Wassif CA, et al. 1H NMR-Linked Metabolomics Analysis of Liver from a Mouse Model of NP-C1 Disease. J Proteome Res 2016; 15: 3511–3527.

Sandhoff K, Harzer K. Gangliosides and Gangliosidoses: Principles of Molecular and Metabolic Pathogenesis. J Neurosci 2013; 33: 10195–10208.

Sango K, Yamanaka S, Hoffmann A, Okuda Y, Grinberg A, Westphal H, et al. Mouse models of Tay–Sachs and Sandhoff diseases differ in neurologic phenotype and ganglioside metabolism. Nat Genet 1995; 2: 170–6.

Schniepp R, Strupp M, Wuehr M, Jahn K, Dieterich M, Brandt T, et al. Acetyl-DL-leucine improves gait variability in patients with cerebellar ataxia-a case series. Cerebellum & ataxias 2016; 3: 8.

Son SM, Park SJ, Lee H, Siddiqi F, Lee JE, Menzies FM, et al. Leucine Signals to mTORC1 via Its Metabolite Acetyl-Coenzyme A. Cell Metab 2018; 0: 1–10.

Strupp M, Teufel J, Habs M, Feuerecker R, Muth C, Van De Warrenburg BP, et al. Effects of acetyl-DL-leucine in patients with cerebellar ataxia: A case series. J Neurol 2013; 260: 2556–2561.

Tighilet B, Leonard J, Bernard-Demanze L, Lacour M. Comparative analysis of pharmacological treatments with N-acetyl-dl-leucine (Tanganil) and its two isomers (N-acetyl-L-leucine and N-acetyl-D-leucine) on vestibular compensation: Behavioral investigation in the cat. Eur J Pharmacol 2015; 769: 342–349.

Vibert N, Vidal P-P. *In vitro* effects of acetyl-DL-leucine (tanganil®) on central vestibular neurons and vestibulo-ocular networks of the guinea-pig. Eur J Neurosci 2001; 13: 735–748.

Te Vruchte D, Speak AO, Wallom KL, Al Eisa N, Smith DA, Hendriksz CJ, et al. Relative acidic compartment volume as a lysosomal storage disorder-associated biomarker. J Clin Invest 2014; 124: 1320–1328.

Wiederschain GY. The metabolic and molecular bases of inherited disease. Biochem 2002; 67: 611.

Williams IM, Wallom K, Smith DA, Eisa N Al, Smith C, Platt FM. Neurobiology of Disease Improved neuroprotection using miglustat, curcumin and ibuprofen as a triple combination therapy in Niemann – Pick disease type C1 mice. Neurobiol Dis 2014; 67: 9–17.

Yanagisawa H, Ishii T, Endo K, Kawakami E, Nagao K, Miyashita T, et al. L-leucine and SPNS1 coordinately ameliorate dysfunction of autophagy in mouse and human Niemann-Pick type C disease. Sci Rep 2017; 7: 1–9.

Yanjanin NM, Vélez JI, Gropman A, King K, Bianconi SE, Conley SK, et al. Linear clinical progression, independent of age of onset, in Niemann-Pick disease, type C. Am J Med Genet Part B Neuropsychiatr Genet 2010; 153B: 132–140.

Yudkoff M. Brain metabolism of branched-chain amino acids. Glia 1997; 21: 92–8.

