## Supplementary tables for "Acetyl-Leucine slows disease progression in lysosomal storage disorders"

### Supplemental Experiment Procedures

#### ADLL and Enantiomers

ADLL (Molekula #73891210), ALL (Sigma Aldrich #441511), ADL (Sigma #A0876) and Miglustat (600 mg/kg/day) (Oxford GlycoSciences) were administered as dry powder in powdered mouse diet (PicoLab Rodent Diet 20/5053, LabDiet) at a dose of 0.1g /kg/ day for *in vivo* studies. Treatment groups for *in vivo* study consisted of at least 5 mice per group.

*In vitro* studies were conducted with each compound dissolved in PBS to generate a 50mM stock solution and diluted to a 1mM working concentration in media (DMEM High Glucose-Gibco #10566016).

#### Mouse behavioural analysis

Gait analysis was performed using CatWalk 10.5 system (Noldus) according to the manufacturer's instructions and five runs were recorded per animal at each time point. The camera was set to 40cm below the walkway, the walkway was approximately 4cm wide, and detection settings set to camera gain 14.05 and green intensity 0.12. Motor function was measured using an accelerating NG Rota Rod for mice (Ugo Basile) starting from 1rpm to 10 rpm, accelerating every 30 seconds by one rpm.

#### Sample preparation

**Western blot analyses.** Western blot analysis was carried out by homogenising half cerebellums (ranging from 15 to 30 mg wet weight) in RIPA buffer (Pierce RIPA Buffer, Thermo Fischer Scientific-89900) with protease/phosphatase inhibitor cocktail (Halt™ Protease and Phosphatase Inhibitor Cocktail-100X, Cat No: 78440) to achieve 50mg/mL w/v. Samples then incubated for 30 mins on ice. They were centrifuged at 13,000 g at 4°C and supernatants were retained for BCA protein assay and western blotting.

**ADP and ATP extraction.** Freshly prepared tissues were homogenized with 3.0 mL of ice-cold phenol–TE (Sigma-77607) and processed with three cycles of 30-s homogenization and 30-s cooling. One mL of homogenate was transferred into tubes containing 200uL of chloroform and 150uL of de-ionized water. The homogenate was thoroughly shaken for 20s and centrifuged at 10,000 × g for 5 min at 4°C. The supernatant was diluted 1000-fold with deionized water, the diluted extract was used for ADP/ATP analyses.

**NAD and NADH extraction.** Tissues were homogenised with a Dounce homogeniser with 400uL of NADH/NAD extraction buffer (Abcam NAD/NADH Assay kit-ab65348), followed by centrifugation for 5 mins at 4°C at 13.000g. Supernatants were collected into a new tube and filtered through 10kD Spin column (Abcam, #ab93349) at 10,000 g at 4°C until the majority of the liquid had passed through (approximately 20 mins).

#### Sample analysis

**Sphingoid Base Measurements.** Sphingoid bases (sphingosine and sphinganine) were extracted from 100uL tissue homogenates in 500uL of chloroform:methanol (1:2v/v) followed by sonication for 10 mins at room temperature. Subsequently, 1M 500-uL sodium chloride, 500 uL chloroform, and 3M 100uL sodium hydroxide were added to the samples, and vortexed every five mins for 15 mins at room temperature. Homogenates were centrifuged (13,000 g for 10 mins) and the lower organic phase retained. Sphingoid bases were purified from the samples by pre-equilibrating SPA-NH2 columns with 2x1mL chloroform followed by sample elution with 3x300ul acetone. The samples were dried under nitrogen. Lipids were resuspended in 50uL pre-warmed ethanol (37°C) and 50 uL OPA labelling solution (1mg OPA/20 uL Ethanol/1uL 2-mercaptoethanol; dilution 1:2000 in 3% boric acid pH 10.5) was added. Samples were kept at room temperature in the dark for 20 mins and vortexed every 10 mins. Samples were buffered with 100uL methanol: 5mM Tris pH 7 (9:1) and centrifuged at 5,000 g for 2 mins. Supernatants (150uL) were loaded onto a reverse phase HPLC. Chromatography was carried out using a mobile phase of 85% acetonitrile/15% $H_2O$ at a flow rate of 1.0ml/min. The orthophthaldehyde-labelled derivatives were monitored at an excitation wavelength of 340nm and an emission wavelength of 450nm. Quantification of trace peak area was carried out using EZChrom Elite software v3.2.1 (http://www.jascoinc.com/ezchrom).

**Glycosphingolipid measurements.** Glycosphingolipids were extracted and measured by NP-HPLC according to published methods (Neville *et al.*, 2004). Briefly, tissue homogenates in chloroform/methanol (C:M) (1:2 v/v) were kept overnight at 4°C. The mixture was centrifuged (3000rpm/10min) and 1mL chloroform and 1mL PBS were added to the supernatant and centrifuged (3000rpm/10min). The lower phase was dried under $N_2$, resuspended in 50uL C:M 1:3 v/v and combined with the upper phase. Subsequently, GSLs were recovered using C18 Isolute columns (100mg) (Biotage) pre-equilibrated with 4x1mL MeOH and 2x1mL $H_2O$. Samples were washed 3x1mL $H_2O$ and eluted with 1mL C:M 98:2, 2x1mL C:M 1:3, 1mL MeOH. The column eluate was dried under $N_2$, resuspended in 100uL C:M 2:1, dried again under $N_2$ and resuspended in ceramide glycanase (CGase) buffer (50mM sodium acetate pH 5.5, 1mg/mL sodium taurodeoxycholate). 50mU CGase was added, and samples incubated at 37°C overnight. Released oligosaccharides were anthranilic acid (2-AA) labelled and purified by mixing with 1mL Acetonitrile: $H_2O$ 97:3 and added to Discovery DPA-6S columns (SUPELCO, # 52625-U) pre-equilibrated with 1mL acetonitrile, 2x1mL $H_2O$, and 2x1mL acetonitrile. Columns were washed with 2x1mL Acetonitrile: $H_2O$ 95:5 v/v, and eluted in 2x 0.75mL $H_2O$. Samples were loaded 30:70 $H_2O$: MeCN (v/v) for normal phase HPLC according to published method (Neville *et al.*, 2004).

**Cholesterol measurements.** Cholesterol was measured with the Amplex Red kit (Molecular Probes) according to the manufacturer's instructions. Briefly, cell and tissue homogenates in 100uL mQ water were Folch extracted and dried down under nitrogen. Cell pellets containing cholesterol were resuspended in 1X Reaction Buffer, and 50uL was loaded for each sample in a flat bottom 96-well plate. The reaction was initiated by adding Working Solution per sample (25% Amplex® Red, 2U/mL HRP stock solution, 2 U/mL cholesterol oxidase stock solution, 0.2 U/mL cholesterol esterase stock solution in 1X Reaction Buffer). Samples were incubated at 37°C for 30 mins and fluorescence was measured in a microplate reader (Optima, BMG Labtech) using excitation in the range of 530–560 nm and emission detection at ~590 nm.

#### Flow cytometry experiments of CHO cells

*In vitro* FACS experiments were used to measure relative acidic compartment volume staining in live cells with LysoTracker™ Green DND-26 (Thermo Fisher-L7526). Cells were incubated with 250nM LysoTracker in PBS for 10 mins at RT, centrifuged at 1200rpm for 10 mins and resuspended in FACS buffer (PBS, 1% BSA, 0.1% sodium azide ($NaN_3$). Wild type and NPC1 null Chinese Hamster Ovary (CHO) cells were cultured as previously described (Cruz *et al.*, 2000). Cells were stained with propidium iodide (20nM in PBS) (Invitrogen-P3566) immediately prior to analysis on the FACS for dead cell discrimination.

Mitochondrial volume was measured using MitoTracker Green (Invitrogen #M7514), and mitochondrial reactive oxygen species was measured with MitoSOX Red (Invitrogen #M36008). Live cells were incubated in media with 50nM MitoTracker Green and 5uM MitoSOX Red for 10 minutes at 37°C. Cells then washed two times with warm HBSS containing 5% FCS. MitoTracker green was detected in the FITC channel and MitoSOX-red dye was detected at PE channel. FACS analysis was performed on 10,000 recorded cells using FACS Canto (Becton Dickinson, (BD)) with BD software.

#### Filipin Staining

Cells were fixed with 4% paraformaldehyde, washed 3 x PBS and incubated with 1.5mg/mL glycine for 10 min to quench autofluorescence. Cells were then incubated with 0.05mg/mL filipin (from Streptomyces filipinensis (Sigma)) diluted in PBS containing 10% FBS and 0.2% Triton X-100 for 2 hours at RT. Cells were then washed 3 x PBS, and coverslips mounted onto microscope slides with DPX, before being visualized with a Leica-SP8 confocal microscope.

#### Immunohistochemistry

Brains were cut parasagitally (20-micron sections) and stored in 30% ethylene glycol, 30% glycerol 0.1M sodium phosphate buffer (pH7.4) at -20°C. Free-floating brain sections in glycerol/ethylene were rinsed 3 x PBS and blocked with 2% goat serum in a 0.3% Triton X-100/PBS at RT for an hour. Sections were stained with primary antibodies; rabbit calbindin (1:2000) and rat CD68 (1:500) in 2% goat serum in 0.3% Triton X-100/PBS for 16 hours at 4°C, followed by washing with 0.3% Triton X-100/PBS, then 3 times with PBS. Secondary antibody staining (Alexa Fluor 594, goat anti-rabbit Red, Alexa Fluor 488, goat anti-rat) was conducted at RT for 2 hours. The antibodies used are summarised in **Supplementary Table 1**. Samples were washed once with 0.3% Triton X-100/PBS, 3 x PBS and manually mounted onto slides.

#### Western Blotting

Homogenates of mouse cerebellum were prepared by Dounce homogenising tissue at 50mg wet weight in 1mL of lysis buffer (1% Igepal CA, 0.5% sodium deoxycholate, 0.1% SDS, and 1% protease-phosphatase inhibitor cocktail) and and protein content was determined using a BCA protein assay (Thermo Fisher #23227) according to the manufacturer's instructions. Samples were resolved using 4-12% SDS tris-glycine (Biorad) gel electrophoresis and transferred to low fluorescent PVDF membrane (Biorad Immunblot Low Fluorescence PVDF paper; #1620262) using a semidry transfer apparatus (Trans-Blot Turbo Transfer System (Biorad; #1704150)). Antibodies used are summarised in **Supplementary Table 1**.

#### ADP/ATP measurements

Measurements of ADP/ATP were made using a kit from Sigma Aldrich (Cat#: MAK135) according to the manufacturer's instructions. The ADP/ATP ratio was calculated with the equation: (RLU-C-RLUB)/RLU-A. 10ul ADP/ATP extractions were loaded in 96 well plate and 90ul ATP reagent (consisting of assay buffer, substrate, co-substrate and ATP enzyme) was added on samples. Luminescence was read after 1 min incubation (RLU-A). Luminescence was recorded again after 10 minutes incubation (RLU-B). 5 ul ADP reagent (ADP enzyme and water) was added on wells immediately after RLU-B recording, and the third luminescence value was read after 1 min incubation (RLU-C).

#### NAD, NADH measurements

NAD, NADH and total NAD (NADt) (NAD+NADH=NADt) were measured with the NAD/NADH assay kit (Abcam-ab65348). Fresh samples were processed for NAD-NADH extraction. Each sample was divided into two: (1) Normal samples and (2) Decomposed samples. Decomposition of each sample was achieved by heating samples at 60°C for 30 mins on a heating block. 30uL extracts were loaded in triplicate into flat bottom clear 96-well plates, for each respective condition: (1) Background, (2) NADt and (3) Decomposed samples (contains only NADH). Background reaction mix (60uL) was loaded into "background wells", and Reaction Mix was loaded into "NADt" and "Decomposed sample" wells. The plate was incubated at room temperature for 5 mins, and 6uL NADH developer solution was added to the "Decomposed sample" wells. Absorbance readings were taken at 30 min intervals over 3hr at 450 nm.

#### Clinical Studies

##### Demographics and statistical analysis of a clinical observational study of adult NPC1 patients treated with ADLL

A cohort of thirteen NPC1 patients participated in this study, comprising 3 females and 10 males from Germany, Saudi Arabia, Sweden, Bulgaria, Slovakia, Turkey and the Czech Republic. Informed consent from all participants was obtained. Twelve of the NPC1 patients were on miglustat therapy (minimum exposure 2.65 years, 1st quartile 3.44, median 4.92, mean 4.87, 3rd quartile 5.80 and maximum 7.98 years). The thirteen NPC1 patients participated in an observational study involving treatment with 5g/day acetyl-DL-leucine (Tanganil™). Three patients had four prospective clinical severity score measurements and ten patients had five measurements taken during the study (NIH, CSS) (Yanjanin *et al.*, 2010). Retrospective data were used to calculate rates of disease progression prior to recruitment to this study. The mean age at the start of acetyl-DL-leucine (Tanganil™) treatment was 26.59 years. The minimum age was 15.44, 1st quartile 20.97, median 26.98 and third quartile 30.79 with maximum age 37.37 years. The time in the observational treatment study was a mean of 0.53, 1st quartile 0.81, median 1, mean 0.95, third quartile 1.13 and maximum 1.21 years.

**Individual-cases of off-label-use in GM2 gangliosidosis patients**

In a prospective observational case series in three patients from Germany with genetically proven GM2 gangliosidoses (2 patients with Tay-Sachs disease, 1 patient with Sandhoff disease) we evaluated the symptomatic effects of ADLL. Patients were treated with 0.1 g per kg body weight per day, for at least four weeks and the clinical effects determined using the Scale for the Assessment and Severity of Ataxia (SARA), the 8-meter walk test (8MWT) and the Montreal Cognitive Assessment (MoCA). Consent from all participants was obtained.

Patient 1:

A 28-years-old male patient with a genetically confirmed diagnosis of Tay-Sachs disease presented with dysarthrophonia, tremor, ataxia of stance and gait, paraparesis, and muscular atrophy. Prior to treatment, the patient required the support of both a caregiver on one side and a wall on the other in order to walk down a corridor. The patient was receiving sodium-valproate (500 mg/d), lithium (450 mg/d), and risperidone (3 mg/d) (medications for clinically diagnosed bipolar disorder), and continued to do so during treatment with ADLL. The patient was also receiving physiotherapy, ergotherapy, and logotherapy twice a week.

Patient 2:

A 32-year-old female patient with a genetically confirmed diagnosis of Tay-Sachs disease presented with ataxia of stance and gait, fine motor impairment, paraparesis of lower extremities and muscular atrophy. She was receiving physiotherapy, ergotherapy and logotherapy and continued to do so during treatment with N-Acetyl-DL-Leucine.

Patient 3:

An 8-year-old male patient suffering from Sandhoff disease, with muscle hypotrophy, epileptic cramps (tonic-clonic, about 10 seconds, self-limiting), disordered ocular movement and anarthria. His daily activities were very limited, being unable to eat, wash, or dress himself. He was receiving levetiracetam (1 g/d), lamotrigine (100 mg/d), omeprazole (20 mg/d), and miglustat (300 mg/d), in addition to physiotherapy, ergotherapy and logotherapy, and continued to do so during treatment with N-Acetyl-DL-Leucine.

**Blinded Video-Rating**

Three international movement disorder experts from Germany (1) and Switzerland (2) evaluated the videos blinded to treatment status and the dates when the recordings were made. The videos were rated based on the Clinical Impression Change in

Severity (CI-CS), as follows: 1 = normal, not at all ill; 2 = borderline ill; 3 = mildly ill; 4 = moderately ill; 5 = markedly ill; 6 = severely ill; 7 = among the most extremely ill patients. After unblinding, their ratings were evaluated regarding the change between the baseline and on medication time point.

#### Statistical methods

##### Murine studies

Differences between mouse experimental groups were identified by 1-way or 2-way analyses of variances (ANOVA); comparisons of groups were made after 2-way ANOVAs (with 95% confidence interval) with Fisher's least significance difference (LSD) test to adjust for multiple comparisons, where necessary. Statistical tests were performed using GraphPad Prism 6 software (La Jolla, CA). For more than three groups, the LSD test is increasingly conservative. This means that if a post-hoc comparison is significant accordingly to LSD then it's likely to be significant for other, possibly more powerful tests, e.g. Benjamini-Hochberg's. Since we are doing a large number of post-hoc comparisons, using LSD can be considered as protective against possibly large discovery rates so it was selected to achieve increased rigour in the analyses.

##### Blinded video rating

Blinded video rating results were calculated with t-test (SPSS). This was statistically significant ($p$=0.0039, **Figure 7d** and **Table 2**).

| *Abv/acronym* | *Antibody type* | *Host* | *Source* | *Catalogue code* | *Dilution* |
|---|---|---|---|---|---|
| Calbindin-D28K | Mab | Rabbit | Swant | CB38a | 1:2000 |
| CD68 | Mab | Rat | Bio-Rad | MCA1957 | 1:500 |
| Alexa Fluor 594 anti rabbit | Secondary Ab | Goat | abcam | ab150080 | 1:1000 |
| Alexa Fluor 488 anti rat | Secondary Ab | Goat | abcam | ab150157 | 1:2000 |
| LC3B | Polyclonal | Rabit | abcam | ab51520 | 1:2000 |
| P62 | Mab | Mouse | abcam | ab56416 | 1:2000 |
| GDH | Mab | Rabit | CellSignalling | D9F7P | 1:1000 |
| pMTOR (ser 2448) | Polyclonal | Rabit | CellSignalling | 2971 | 1:500 |
| mTOR | Polyclonal | Rabit | CellSignalling | 2972 | 1:500 |
| PDH Complex | Polyclonal | Mouse | abcam | ab110416 | 1:1000 |
| pPDH (s293) | Polyclonal | Rabit | abcam | ab92696 | 1:500 |
| SOD1 | Polyclonal | Rabit | abcam | ab13498 | 1:1500 |
| SOD2 | Polyclonal | Rabit | abcam | ab56416 | 1:1500 |
| PDP1 | Polyclonal | Rabit | abcam | ab228578 | 1:1000 |
| PDK1 | Mab | Mouse | abcam | ab110025 | 1:1000 |
| PDK2 | Mab | Rabit | abcam | ab68164 | 1:1000 |
| PDK4 | Mab | Rabbit | abcam | ab214938 | 1:1000 |
| BCKADH-A | Polyclonal | Rabbit | abcam | ab90691 | 1:1000 |
| pBCKADH-A | Polyclonal | Rabbit | abcam | ab200577 | 1:1000 |
| LDHB | Mab | Mouse | abcam | ab85319 | 1:1500 |
| PGC-1 alpha | Mab | Mouse | Merck Millipore | ST1202 | 1:1500 |
| Anti-Mouse 800CW IgG (H + L) | Secondary Ab | Goat | Licor-IRDye | 925-32210 | 1:10,000 |
| Anti-Rabbit 680RD IgG (H + L) | Secondary Ab | Goat | Licor-IRDye | 925-68071 | 1:10,000 |
| HRP conjugated beta actin | Mab | Mouse | Invitrogen | MA5-15739-HRP | 1:15,000 |
| Pierce ECL Substrate Kit | - | - | Thermo Fisher | 32106 | - |

**Supplementary Table 1.** List of antibodies /reagent used for immunohistochemistry and western blotting.
