## Supplementary figures and images for "Acetyl-Leucine slows disease progression in lysosomal storage disorders"

### Supplemenary figures

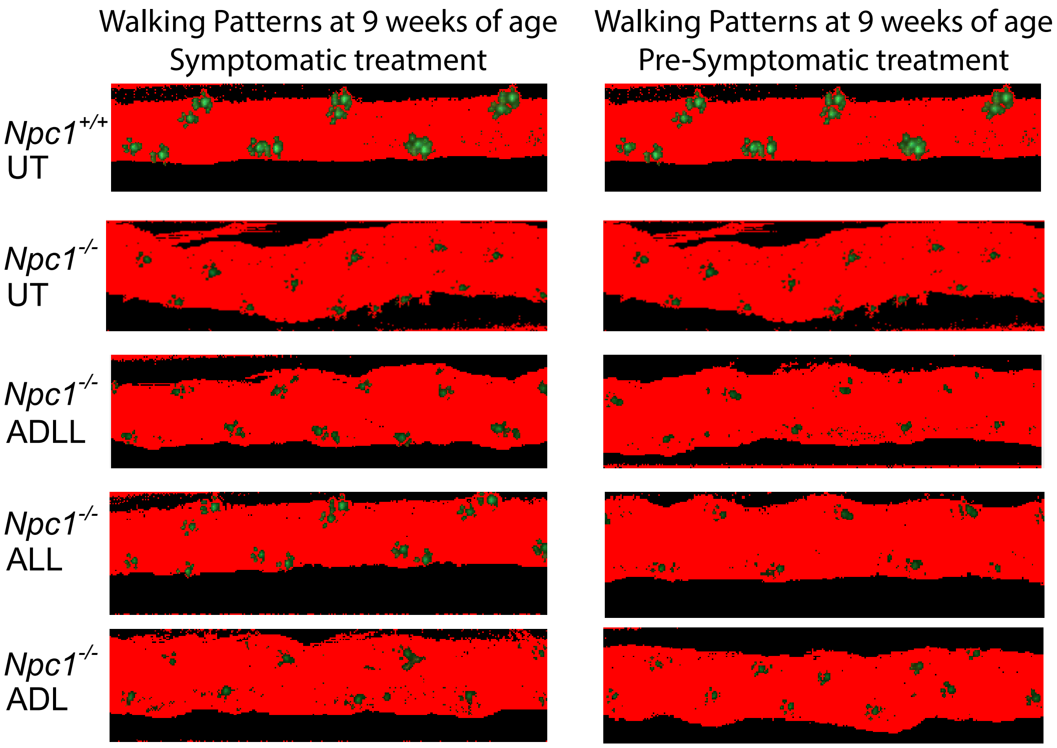


**Supplementary Figure 1.**

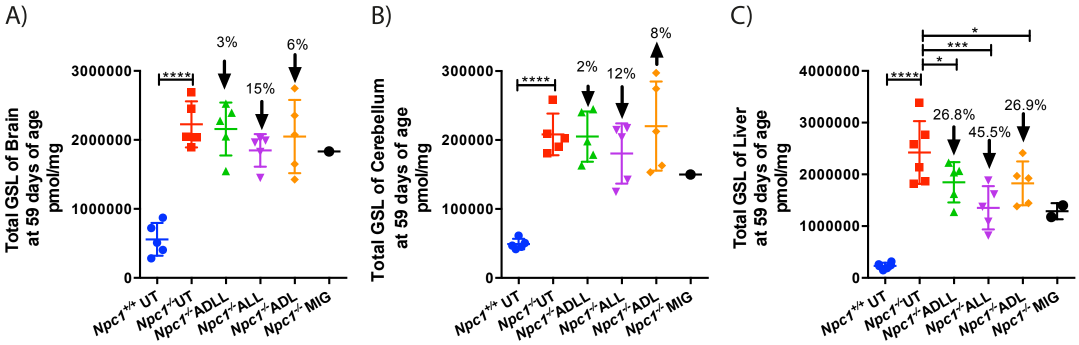


**Supplementary Figure 2.**

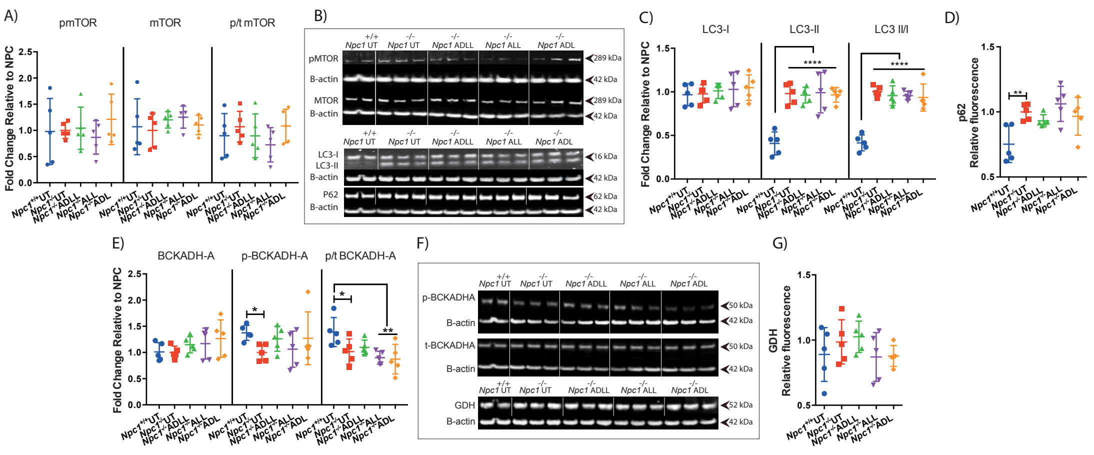


#### Supplementary Figure 3.
